# A network-based deep learning model integrating subclonal architecture for therapy response prediction in cancer

**DOI:** 10.64898/2026.03.14.711567

**Authors:** Sungnam Kim, Doyeon Ha, A-reum Nam, Sehyoung Cheong, Juhun Lee, Sanguk Kim, Solip Park

## Abstract

Predicting treatment response remains challenging in oncology, particularly given the growing diversity of therapeutic options. Despite efforts using gene expression signatures, or integrative multi-omics frameworks, robust and interpretable biomarkers remain limited. We present SubNetDL, a deep learning framework that integrates subclonal mutation profiles and protein–protein interaction networks via network propagation. Unlike condition-specific approaches, SubNetDL leverages somatic mutations alone and is applicable across diverse cancer types and treatment modalities. Applied to ten TCGA cancer–drug combinations, SubNetDL achieved consistently strong performance (median AUROC = 0.74) and successfully generalized to two independent immunotherapy datasets (median AUROC = 0.77). Importantly, it identified candidate biomarker genes with treatment-specific relevance. SubNetDL prioritized genes that were not central in the network, highlighting its ability to capture context-specific patterns beyond traditional metrics. In conclusion, our approach offers a robust and interpretable framework for identifying predictive biomarkers and stratifying patients based on mutation profiles and network context.

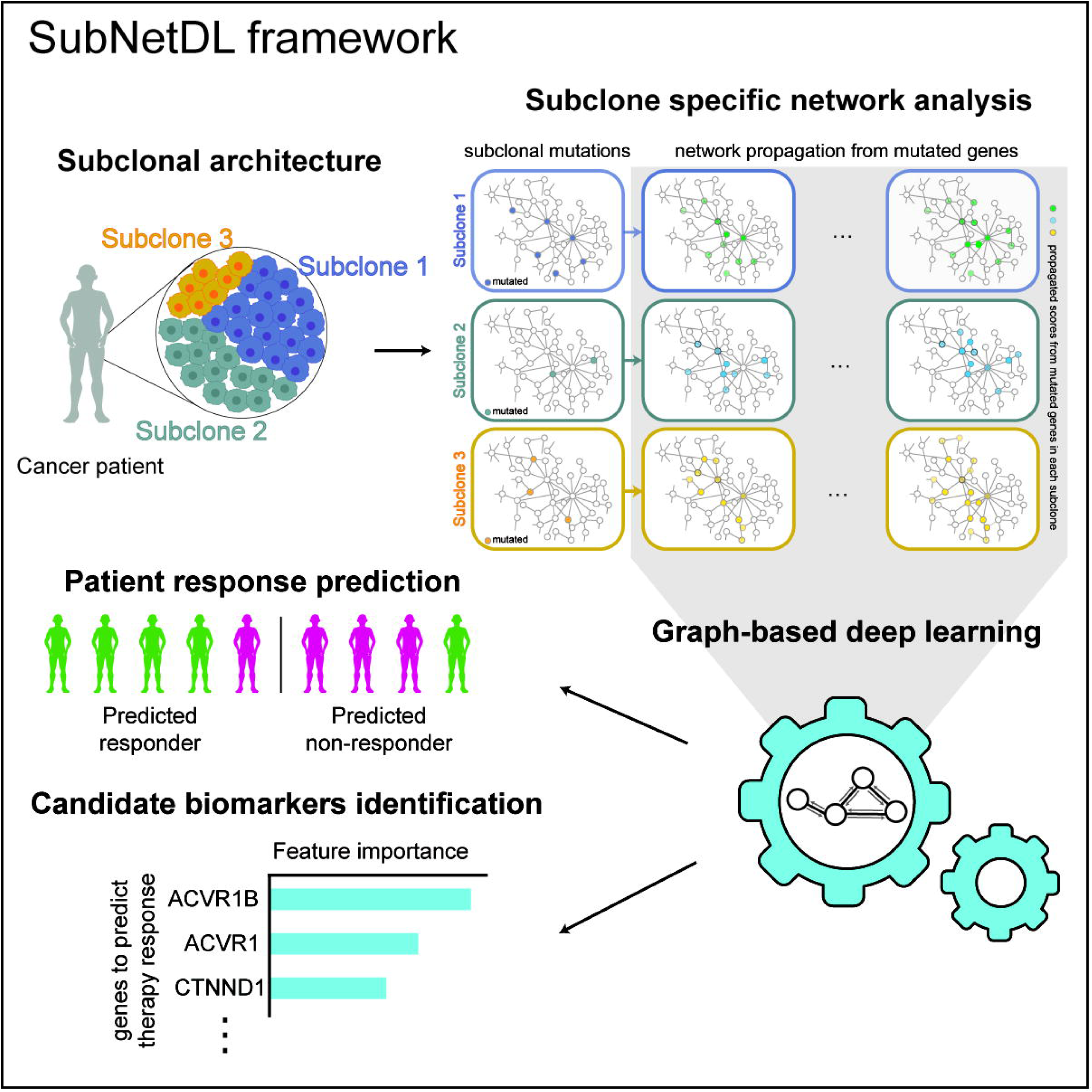

**Motivation:** Intratumoral heterogeneity is a fundamental driver of therapeutic resistance, yet most predictive models rely on aggregate mutational burdens or static gene expression signatures, overlooking the subclonal dynamics that shape treatment outcomes. While network biology offers a functional lens to interpret genomic alterations, a framework that explicitly bridges subclonal architecture with system-level molecular interactions has been lacking. To address this, we developed SubNetDL, a deep learning framework that integrates patient-specific subclonal profiles with protein-protein interaction networks. By leveraging only somatic mutation data, SubNetDL captures the functional convergence of subclonal evolution, providing a robust and interpretable platform for patient stratification and biomarker discovery across diverse oncological contexts.

## Introduction

Over the past two decades, anti-cancer therapies—ranging from chemotherapy to targeted therapy, immunotherapy, and combinatorial or multimodal regimens—have led to significant survival gains in many cancer types^1–3^. However, predicting therapeutic response remains a central challenge in precision oncology^4^. Despite the discovery of various biomarkers, such as gene expression signatures, tumor mutational burden (TMB), and developing drug-specific machine learning (ML) classifiers, current predictors often exhibit limited generalizability across patients, cancer types, and treatment modalities^5–8^.

This is exemplified by immunotherapy, where only three U.S. Food and Drug Administration (FDA)-approved biomarkers—programmed cell death ligand-1 (PD-1) expression, TMB, and microsatellite instability (MSI) status—are currently in clinical use^9,10^. Even so, their predictive performance varies markedly by tumor type and patient context, underscoring the broader issue of inconsistent biomarker reliability^5^. These limitations largely stem from the intrinsic molecular heterogeneity of tumors, which encompasses not only diverse mutation profiles but also the dynamic subclonal architecture and functional contexts in which these alterations operate^11,12^.

While many existing models have demonstrated utility in predicting treatment response, they often overlook intratumoral heterogeneity. This heterogeneity poses a major confounding factor, as different subclones within a tumor may exhibit varying sensitivities to therapy—some being susceptible, while others are resistant. The presence of specific mutations within certain subclones can drive therapeutic resistance, as exemplified by *EGFR*-mutated non-small lung cancer, where secondary T790M mutations arise under tyrosine kinase inhibition^13,14^; by melanoma, where *BRAF* inhibitor resistance often emerges through subclonal alterations in MAPK pathway components^15^. Beyond resistance mechanisms, increasing evidence supports a direct link between subclonal diversity and clinical outcomes across cancer types. For instance, subclonal diversification has been associated with poor prognosis in chronic lymphocytic leukemia, ovarian, breast, and lung cancers, among others^16–21^. Subclonal somatic alterations—such as CNAs and mutational drivers—correlate with reduced survival and higher risk of relapse, underscoring the biological and clinical significance of subclone-level analyses^22^. These findings highlight the growing recognition that incorporating clonal architecture into predictive models is essential for capturing the biological complexity of tumors and improving clinical outcome predictions^23,24^.

In parallel, network biology approaches have been utilized to capture the biological relevance embedded within tumor heterogeneity. For example, distinct genetic variants may perform similar biological functions, which can be revealed through network-based analyses^25^. In protein-protein interaction (PPI) networks, functionally related genes—such as those within protein complexes or shared signaling pathways—tend to form clusters, as edges in the network represent functional associations^26^. Accordingly, network propagation approaches have been successfully used to identify patient subtypes across heterogeneous cancer populations by capturing the functional convergence of mutations on shared pathways^27^. Building upon this framework, our previous work successfully demonstrated that network-based diffusion of somatic mutations could predict response to immune checkpoint blockade across cancer types^28–31^.

Building on these observations, we present a deep learning framework that integrates subclonal structure and network-based propagation to predict therapeutic response. By leveraging both the clonal composition of tumor mutations and their diffusion across PPI networks, our model captures complex, tumor-specific determinants of drug sensitivity. We evaluated its performance across ten cancer–drug pairs from TCGA and further validated its applicability in immunotherapy settings, demonstrating consistent predictive accuracy and mechanistic interpretability. Overall, these findings highlight the importance of network-level convergence and subclonal diversity in shaping treatment response.

## Results

### A subclonal architecture-informed network-based deep learning framework for predicting anti-cancer therapy response

We developed SubNetDL, a deep learning framework to predict anti-cancer drug response by integrating tumor subclonality with network-based propagation features. To capture the predictive value of tumor clonality in the context of molecular interaction network, the framework was made up of three key components: i) subclonal inference, ii) network propagation using subclone-specific mutational profiles, and iii) network-based deep learning for therapeutic response prediction. (**Fig. 1A**).

**Figure 1.**
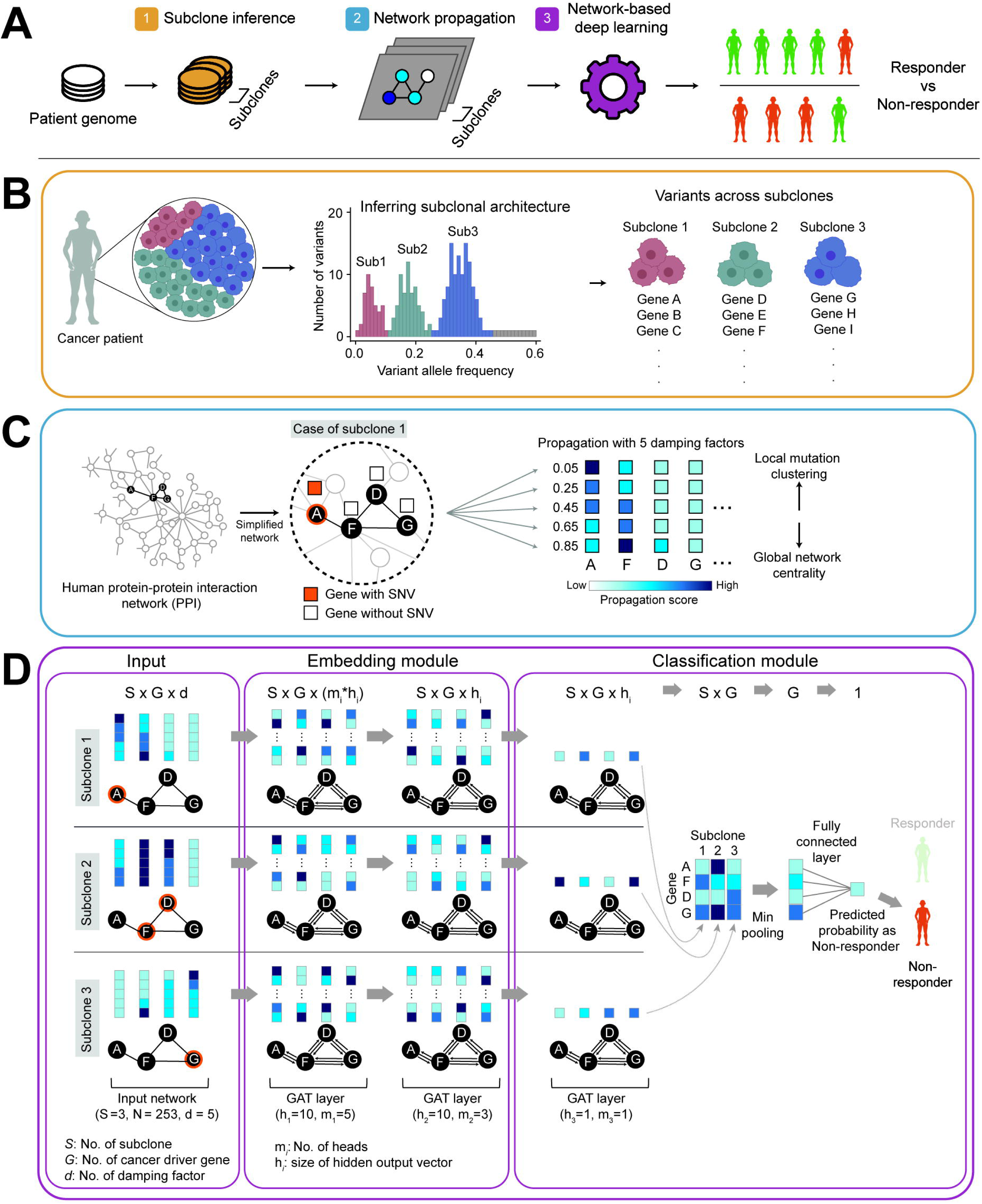
Overview of SubNetDL, a network-based deep learning framework for predicting cancer drug response from patient-specific somatic mutations. (**A**) Overview of the pipeline. Patient tumor genomes are analyzed to identify somatic mutations, which are then grouped into subclones based on their variant allele frequency (VAF) profiles. For each subclone, network propagation is applied to a protein–protein interaction (PPI) network, which serve as input to a deep learning model. The model integrates features across all subclones to predict whether the patient is likely to respond to a given anti-cancer drug. (**B**) Subclone inference. Mutations with similar VAFs are clustered to define subclones, capturing tumor heterogeneity. **(C)** Network propagation. For each subclone, a subclone-specific network is constructed by applying network propagation on the full PPI graph, where mutated genes (proteins) are initialized with a value of 1 and non-mutated genes with 0. Propagation is performed using the PageRank algorithm with five different damping factors (0.05, 0.25, 0.45, 0.65, 0.85), resulting in a five-dimensional propagation score for each gene. Lower values emphasized the influence of proximal nodes, capturing local mutation clustering, while higher values allowed broader signal diffusion across the network, reflecting global topological context. To focus on cancer-relevant signals, the propagated scores were restricted to 253 curated driver genes; within this subset, edges were reweighted based on shortest path distances to construct a simplified, proximity-aware network. (**D**) Network-based deep learning model. Each subclone’s feature matrix is processed independently through a Graph Attention Network (GAT) comprising two multi-head layers (embedding module) followed by a single-head GAT layer, min-pooling, and a fully connected classifier (classification module). The final output is a binary prediction indicating whether the patient is a responder or non-responder. In the illustrative example, three subclones (*S*=3) and four genes (*n*=4) are shown.

The framework started with inferred patient-specific subclonal architecture to account for intra-tumoral heterogeneity using SciClone^32^, which clusters somatic mutations based on variant allele frequency (VAF) distributions (**Fig. 1B**). Based on the inferred architecture, mutations were assigned to their respective subclones, yielding subclone-specific mutation profiles. This process identified between one and six subclones per tumor, providing a stratified view of somatic alterations along the clonal hierarchy.

SubNetDL embeds subclonal mutational landscapes into network topology-aware representations, effectively linking molecular heterogeneity to system-level interaction features. To model the network-level effects of subclonal mutations, we constructed subclone-specific networks by duplicating the global STRING^33^ PPI network (with confidence score > 700), comprising 14,278 nodes and 255,825 edges. Each patient therefore has as many PPI networks as the number of detected subclones (typically ranging from 1 to 6 as inferred by SciClone^32^; average = 1.97 and median = 2.0). Mutation-informed network propagation was performed by assigning binary seed labels (1 = mutated, 0 = not mutated) to the nodes. To capture both local and global interaction contexts, we applied five damping factors (α = 0.05, 0.25, 0.45, 0.65, 0.85; **Supplementary Figure 1A and B**), enabling multiscale propagation of mutational signals through the network (**Fig. 1C**).

SubNetDL employed a Graph Attention Network (GAT) architecture to encode each subclone-specific graph into high-dimensional embeddings^34^. GAT extends self-attention to graph domains by learning edge-wise attention weights, which enables the model to prioritize biologically relevant neighbors during message passing. Multi-head GAT layers were used to capture diverse neighborhood contexts, followed by pooling and fully connected layers to integrate signals across subclones and generate binary predictions of drug response (**Fig. 1D**). To constrain the feature space and enhance interpretability, we pruned the network to include 253 high-confidence cancer driver genes (selected from a curated set of 299^35^) that overlapped with STRING nodes. These genes formed the backbone of the diffusion framework, enabling the modeling of cancer-relevant interaction topologies while reducing complexity (**See, Method**). These components define the architecture of SubNetDL, which we next benchmarked across ten cancer–drug pairs to evaluate its predictive performance.

### SubNetDL exhibits robust predictive performance across cancer–drug cohorts

SubNetDL achieved robust predictive performance across ten cancer–drug pairs (median area under the receiver operating characteristic curve [AUROC]:0.74, range from 0.56 to 0.94) (**Fig. 2A)**. The model was evaluated on a held-out test set following an 8:1:1 stratified split to preserve class balance. Model hyperparameters were tuned on the validation set.

**Figure 2.**
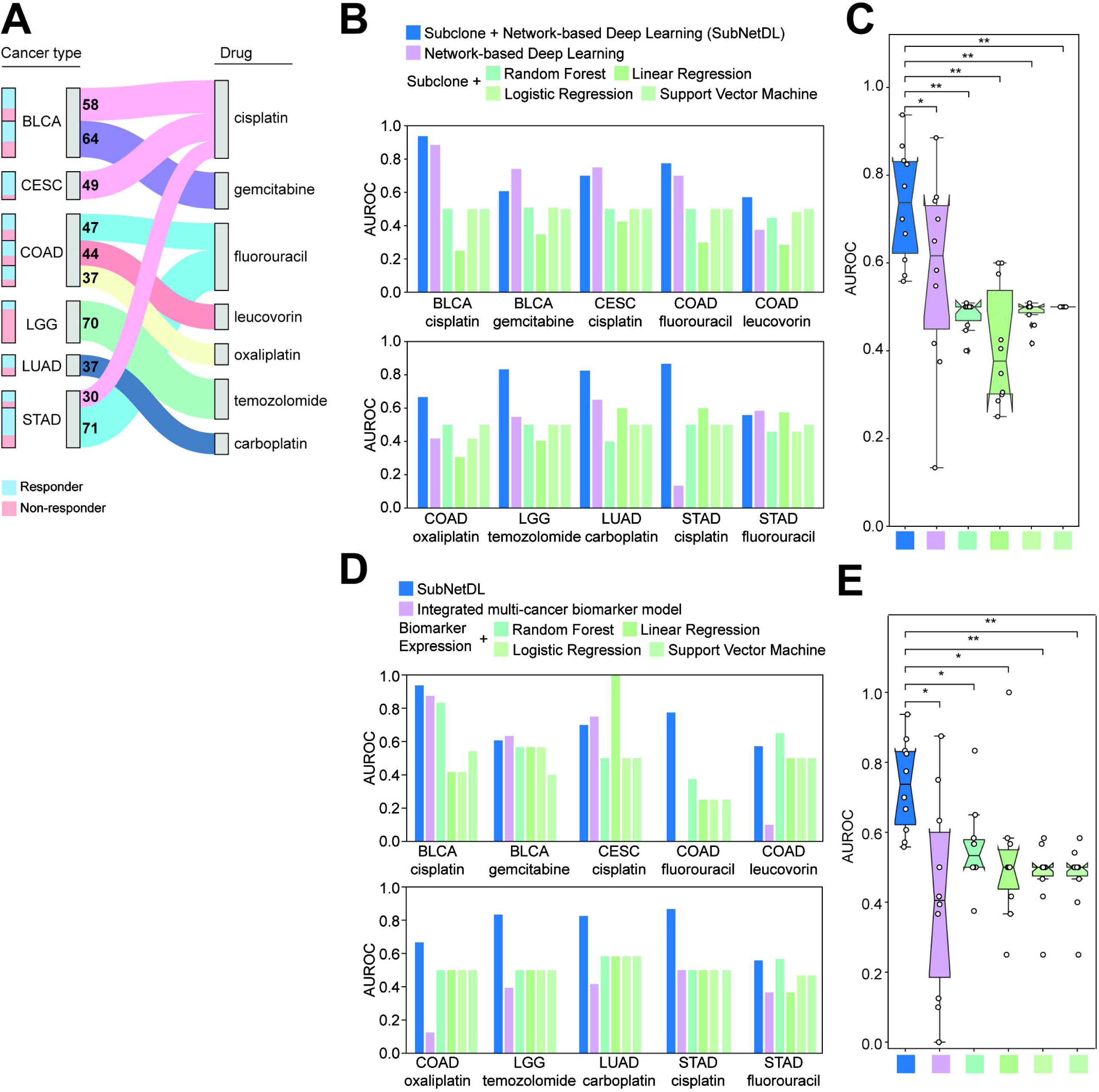
Predictive performance of subclonality-integrated network-based deep learning models across ten cancer type–drug treatment pairs. (**A**) Alluvial plot summarizing the distribution of ten selected cancer–drug pairs used for evaluation. Each pair includes the cancer type, treatment, and response outcome (blue: responders, pink: non-responders), with sample sizes indicated. Cancer–drug pairs were selected based on sample size (*n* > 30) and a balanced distribution of responders and non-responders, resulting in ten pairs from six cancer types: endocervical adenocarcinoma (CESC), lung squamous cell carcinoma (LUSC), colon adenocarcinoma (COAD), stomach adenocarcinoma (STAD), lung adenocarcinoma (LUAD), bladder urothelial carcinoma (BLCA), and brain lower grade glioma (LGG), and seven distinct treatments. (**B**) AUROC scores for drug response prediction across the ten pairs using six models: (i) subclonality-integrated network-based deep learning model (blue; SubNetDL), (ii) network-based model without subclonality (pink), and (iii) four subclonality-based machine learning models using random forest, linear regression, logistic regression, and support vector machine (green). Models were evaluated on a held-out test set using an 8:1:1 split for training, validation, and testing. (**C**) Boxplot summarizing AUROC scores across the ten cancer–drug pairs for each of the six models. Each point represents the AUROC from a single cancer–drug pair. Boxes represent the interquartile range (IQR), center lines denote the median, and whiskers extend to 1.5×IQR. Statistical comparisons between models were performed using paired two-sided Wilcoxon signed-rank tests. *P*-values < 0.05 and < 0.005 are indicated by * and **, respectively. (**D**) Comparison of SubNetDL performance with combined expression-based biomarkers previously reported to predict drug response in cancer patients, using AUROC as the evaluation metric. (**E**) Boxplot summarizing AUROC scores between the ten cancer–drug pairs and expression-based biomarkers with the highest AUROC across cancer types. Boxes represent the interquartile range (IQR), center lines denote the median, and whiskers extend to 1.5×IQR. Statistical comparisons between models were performed using paired two-sided Wilcoxon signed-rank tests. *P*-values < 0.05 and < 0.005 are indicated by * and **, respectively.

The predictive performances of SubNetDL were robustly evaluated using ten cancer-drug pairs. Specifically, we compiled a dataset of 507 patients treated with anti-cancer drugs across six solid cancer types from The Cancer Genome Atlas (TCGA)^36^, including bladder urothelial carcinoma (BLCA; *n* = 122) cervical squamous cell carcinoma (CESC; *n* = 49), colon adenocarcinoma (COAD; *n* = 128), stomach adenocarcinoma (STAD; *n* = 101), lung adenocarcinoma (LUAD; *n* = 37), and brain lower grade glioma (LGG; *n* = 70) (**Fig. 2A**). Each patient received a single drug treatment, though multiple drugs were tested within the same cancer type (e.g., COAD patients received fluorouracil, leucovorin, or oxaliplatin). To ensure statistical robustness, we retained only drug–cancer pairs with at least 30 samples (See, **Methods**). Across most pairs, the mean proportion of responders exceeded that of non-responders (mean: 65.0%, range: 56.25–83.67%), except for LGG, where the proportions were reversed (responders: 20.0%, non-responders: 80.0%).

Individual model components– subclonal inference and network-based deep learning based on propagated subclone profiles – contributed to the best predictive performances. To test this, we compared SubNetDL with two ablation settings: one excluding subclonality-informed features (or subclonal architecture features), and another replacing the deep learning module with conventional machine learning models. In the subclonality-excluded setting, all functional exonic SNVs were aggregated into a single mutation profile per patient without explicit clonal decomposition, and diffusion scores were computed from this unified mutation set. Compared to this baseline, SubNetDL achieved higher median predictive performance (0.74 vs. 0.62), with a marginally significant difference (**Fig. 2B** and **C**, labeled “Network-based Deep Learning”, Wilcoxon signed-rank test, *P* < 0.05). It also outperformed all classical machine learning models trained solely on subclonality-derived features: random forest (AUROC=0.48), logistic regression (0.49), linear regression (0.41), and support vector machine (0.50), with all comparisons statistically significant (*P* < 0.05) (**Fig. 2B** and **C**). Similar trends were observed across other performance metrics—including accuracy, area under the precision-recall curve (AUPRC) and F1 score (the harmonic mean of precision and recall) —where SubNetDL generally outperformed both the subclonality-ablated and classical ML models (**Supplementary Figure 1C-F**), with statistical significance in nearly all cases (*P* < 0.05, Wilcoxon signed-rank test), except for AUPRC. Although SubNetDL achieved a higher median value (0.86 vs. 0.77), the difference was maginally significant (*P* = 0.19). These results highlight the added benefit of jointly modelling subclonal structure and leveraging network-based deep learning for drug response prediction. In contrast to classical machine learning models trained solely on binary mutation status of individual genes—without incorporating subclonality-informed features or deep learning—SubNetDL showed significantly superior performance (median AUROC: 0.72 vs. ≤ 0.42 for all classical models, *P* < 0.05, Wilcoxon signed-rank test; **Supplementary Figure 2A** and **B**). These results underscore the added benefit of jointly modeling subclonal architecture and leveraging network-based deep learning for drug response prediction.

To assess whether SubNetDL predictions translate into clinically meaningful survival differences beyond binary response labels, we performed Kaplan–Meier survival analyses with log-rank testing and estimated hazard ratios using Cox proportional hazards models. Significant survival stratification was observed in at least one endpoint in 7 of 10 cohorts. Specifically, overall survival (OS) reached statistical significance in 4 of 10 cohorts, while progression-free interval (PFI) was significant in 6 of 10 cohorts. These findings indicate that SubNetDL prediction scores capture clinically relevant survival dynamics beyond binary classification metrics (**Supplementary Figure 3A**).

SubNetDL achieved robustly higher predictive performance than gene expression–based classifiers in diverse drug–cancer contexts. We benchmarked SubNetDL against established gene expression–based biomarkers of drug response (**Fig. 2D** and **E**; **Supplementary Figure 2C** and **D**). To reduce potential bias from single-marker analyses, these biomarker sets (4 - 18 genes per combination) were also evaluated within multi-gene machine learning classifiers. Across both single-gene and multi-gene settings, SubNetDL consistently achieved superior or comparable performance, underscoring the robustness of its mutation-informed framework (**Fig. 2D** and **E**; *P* < 0.05 with a Wilcoxon signed-rank test). Considering the heterogeneous cellular composition of tumour samples, expression-based biomarkers can be influenced by microenvironmental variability, potentially limiting their stability. Conversely, SubNetDL leverages subclonality-informed somatic mutation–derived network features, which are comparatively more robust to batch effects and tissue processing variability.

To further contextualize performance relative to recently developed deep learning approaches for drug response prediction, we benchmarked SubNetDL against SubCDR^37^ and NIHGCN^38^ using identical TCGA cancer–drug cohorts and patient-level data splits. SubNetDL achieved a median AUROC of 0.73 (range = 0.56–0.94), compared to 0.67 for SubCDR (range = 0.50–0.80; *P* = 0.28) and 0.68 for NIHGCN (range = 0.50–1.00; *P* = 0.49; **Supplementary Figure 3B and C**). These results indicate that SubNetDL achieves predictive performance comparable to existing state-of-the-art approaches under a controlled evaluation framework.

### SubNetDL uncovers tumor-specific, not drug-specific, response patterns

The most informative genes prioritized by SubNetDL were both essential and sufficient for the predictive performances of the model **(Fig. 2A** and **B)**. Building on the strong predictive performance of SubNetDL, we next aimed to explore the molecular underpinnings of its decision-making process. To this end, we examined gene-level importance scores derived from attention weights in the final layer of the model’s graph attention network (GAT)^34^, aiming to identify the most informative molecular features contributing to prediction across cancer types and treatments. To test whether a reduced subset of top-ranked features could approximate the predictive capacity of the entire model, we trained simplified versions of SubNetDL using only a small fraction of highly ranked genes (based on their importance scores; ranging from 2.5% to 10%) (**Fig. 3A**; see **Methods**).

**Figure 3.**
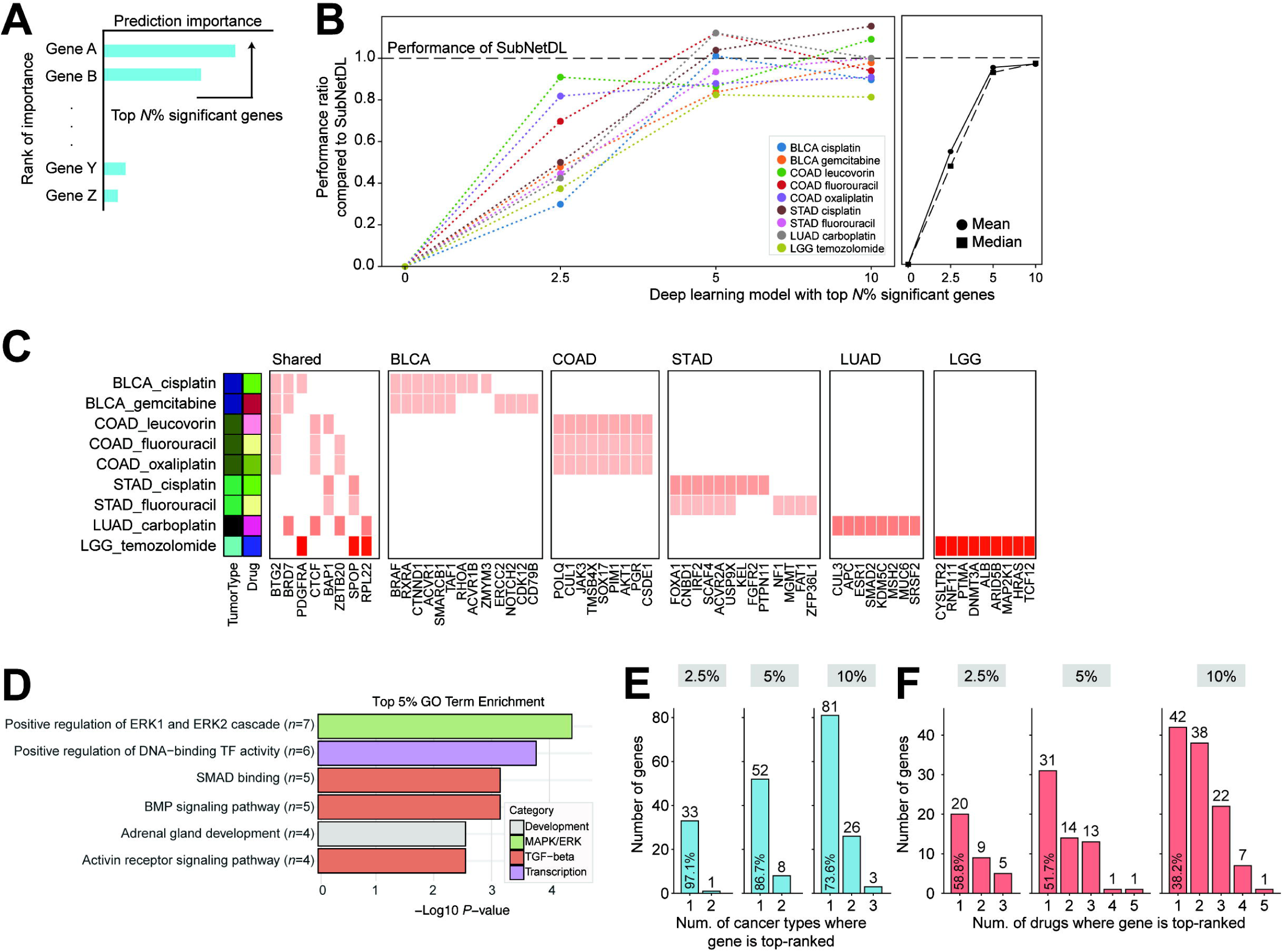
Identification and characterization of significant genes contributing to drug response prediction by SubNetDL. (**A**) Schematic of gene selection process. For each cancer type–drug pair, gene-level importance was quantified using attention scores from the trained SubNetDL model. Genes were ranked based on their importance, and the top *N*% were defined as significant. (**B**) Model performance using only the top *N* % significant genes. A modified version of SubNetDL was constructed using only the selected genes and their corresponding PPI subnetworks as input. AUROC scores were computed on the test set using an 8:1:1 train/validation/test split. The CESC–cisplatin pair was excluded due to the absence of significant genes. (**C**) Heatmap of the top 5% significant genes across all cancer type–drug pairs. Color intensity indicates the relative rank of each gene within a given pair (darker red = higher importance). Genes with consistently high ranks across multiple contexts are positioned toward the left. (**D**) Functional characterization of significant genes. Enrichment analysis was performed using the top-ranked genes to identify overrepresented pathways and biological processes relevant to drug response. (**E**) Histogram showing how many cancer types each significant gene was identified in. Genes appearing in multiple drug treatments within the same cancer type were counted once per cancer type. (**F**) Histogram showing the number of distinct drugs each significant gene was identified for, across all cancer types.

The model trained using important genes prioritized by SubNetDL showed a substantial proportion of the original performance. When restricted to the top 5% of genes—typically corresponding to 11-12 genes per pair—the model maintained a median predictive performance ratio of 0.93 compared to the full version. The inclusion of the top 10% of genes further enhanced the performance, reaching near equivalence with the original model (median ratio: 0.98) (**Fig. 3B**). These results demonstrate that a relatively compact set of genes—those deemed most informative by the model—captures a large fraction of the discriminative signal, allowing for biologically interpretable modeling without sacrificing accuracy.

The most informative genes prioritized by SubNetDL converge on shared biological processes. We performed Gene Ontology (GO) enrichment analysis on the top 5% ranked genes across all cancer type–treatment pairs (**Fig. 3C** and **D)**. This analysis revealed a notable enrichment of TGF-β signaling–related biological processes, including SMAD binding (*P* = 1.42E-05 by Fisher’s test), BMP signaling pathway (*P* = 1.25E-04), and activin receptor signaling pathway (*P* = 4.85E-05). These GO terms were significantly enriched in the top 5% gene group (*versus* human gene background) but not observed when the remaining 95% were tested using the same enrichment framework, emphasizing their selective relevance (**Supplementary Figure 4A**). Importantly, these results were not confined to any single cancer type or treatment, but rather observed across a diverse set of contexts—suggesting that TGF-β signaling represents a shared functional axis recurrently activated in chemoresistant tumors^39^. TGF-β signaling, known for its dual role in tumor suppression and progression, has been implicated in epithelial-to-mesenchymal transition (EMT), immune evasion, and drug resistance-hallmarks of aggressive and treatment-refractory tumors. The consistent emergence of TGF-β–associated terms across multiple model instances suggests that SubNetDL, despite being trained on heterogeneous molecular input, converges upon a biologically coherent representation of drug response. This highlights the model’s capacity to uncover interpretable and mechanistically grounded features beyond mere statistical association.

Although shared biological themes such as TGF-β signaling emerged across cohorts, predictive modeling remained strongly cancer-type–dependent. To directly assess whether common pan-cancer predictors could be identified, we retrained the model in a pooled pan-cancer setting using cisplatin-treated patients across BLCA, CESC, and STAD. While cancer-type–specific models achieved high predictive performance (median AUROC = 0.87), the pooled pan-cancer model showed a marked reduction in performance (AUROC = 0.65; **Supplementary Figure 4B**). These findings indicate that while recurrent pathways may be shared, predictive structure is shaped by tumor-specific molecular contexts rather than a pooled multi-cancer setting.

To further dissect whether shared pathway-level signals reflected common gene-level determinants or context-specific drivers, we examined whether the underlying genes driving these signals were recurrent across cancer types or specific to individual contexts. Although many high-scoring genes were unique to individual cancer–drug contexts, we identified a small group of genes that were consistently prioritized across at least two cancer types at the 5% importance threshold (**Fig. 3C**). This recurrent set included *BTG2*, *BRD7*, *PDGFRA*, *CTCF*, *BAP1*, *ZBTB20*, *SPOP*, and *RPL22*. While functionally diverse, many of these genes play central roles in transcriptional regulation, cell cycle control, or protein degradation pathways^40–42^, and several of these genes have been previously implicated in chemoresistance or are recurrently altered in cancer, with *PDGFRA* and *BAP1* also serving clinically relevant biomarkers in specific cancer types^43,44^. Their repeated prioritization suggests that SubNetDL may be capturing biologically relevant determinants of drug response.

Simultaneously, a comprehensive examination of gene-level prioritization revealed that SubNetDL’s most influential features are largely cancer-type–specific (**Fig. 3E**). The overlap analysis of top-ranked genes revealed that most prioritized genes were unique to individual cancer types—97.1%, 86.7%, and 73.6% at 2.5%, 5%, and 10% thresholds, respectively. Conversely, a notable proportion of genes were shared across different treatments within the same cancer type (41.2%, 48.3%, and 61.8%; **Fig. 3F**). This pattern suggests that while a small set of generalizable, biologically interpretable features such as TGF-β signaling consistently emerges, the majority of SubNetDL’s predictive capacity derives from tumor-specific molecular landscapes that are fine-tuned to each cancer type.

### Validation of SubNetDL in external immunotherapy datasets

SubNetDL generalizes well across independent clinical and preclinical cohorts. Given the strong performance of SubNetDL across TCGA-derived cancer–drug pairs, we next evaluated its generality in independent settings. We first assessed its performance in external immunotherapy-treated patient cohorts and subsequently examined its applicability in independent preclinical cell line datasets. Accordingly, we searched for publicly available immunotherapy cohorts with matched somatic mutation profiles from cBioPortal and collected two eligible cancer cohort datasets from Memorial Sloan Kettering Cancer Center: (i) 242 advanced Non-Small Cell Lung Cancer (NSCLC) patients treated with PD-(L)1 blockade (MSK_2020), including 60 responders and 182 non-responders; and (ii) 240 NSCLC patients treated with anti–PD-(L)1 agents (MSK_2018), with 43 responders and 197 non-responders^45,46^ (**Fig. 4A**). Both datasets were profiled using the MSK-IMPACT panel, which targets 341–468 genes, including somatic mutations and fusions. Given the moderate sample size, we applied 10-fold cross-validation for the model evaluation. While SubNetDL was trained and tested using a fixed 9:1 split, cross-validation enabled more robust performance assessment in these independent cohorts.

**Figure 4.**
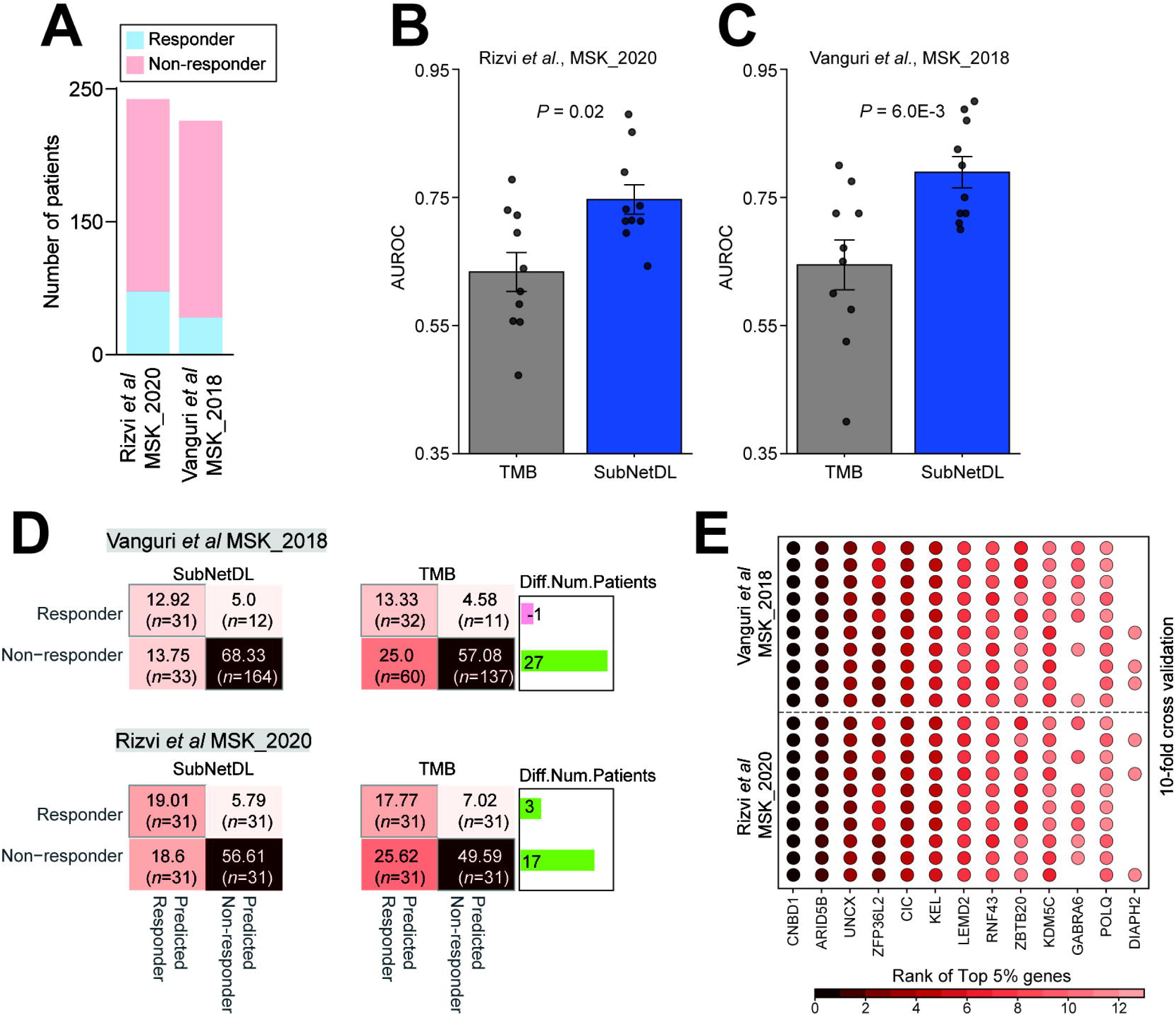
Validation of SubNetDL on independent immunotherapy cohorts (**A**) Overview of immune checkpoint inhibitor-treated NSCLC patient cohorts analyzed using SubNetDL. (**B-C**) Predictive performance (AUROC) of SubNetDL and Tumor Mutation Burden (TMB) in two NSCLC immunotherapy cohorts (lung_msk_mind_2020 and nsclc_pd1_msk_2018). Statistical significance was evaluated using a paired, two-sided Wilcoxon signed-rank test. **(D)** Confusion matrices showing classification performance of SubNetDL and TMB in distinguishing responders vs. non-responders, along with the number of assigned patients in each category. (**E**) Top 5% of genes ranked by significance across cross-validation folds in each cohort. Genes with higher ranks within folds are shown in deeper red, and those with consistently high average ranks across folds are positioned to the left.

SubNetDL achieved strong predictive performance in both cohorts (mean AUROC: 0.75 for MSK_2020; 0.79 for MSK_2018) and significantly outperformed TMB, a widely used biomarker for immunotherapy response (*P* = 0.02; *P* = 6.0E-03 by Wilcoxon signed-rank test, respectively, **Fig. 4B** and **C**). To better understand the performance gain of SubNetDL over TMB, we examined prediction discrepancies across cohorts. SubNetDL more accurately identified both responders and non-responders compared to TMB. In particular, it reduced the number of false positives—patients incorrectly predicted as responders—thereby increasing the true negative rate. Specifically, 27 and 17 patients were correctly reclassified by SubNetDL as non-responders in the MSK_2020 and MSK_2018 cohorts, respectively (**Fig. 4D**). Taken together, these findings indicate that SubNetDL may overcome a key shortcoming of TMB by filtering out likely false positives and providing a more refined framework for stratifying responders to immunotherapy.

In addition to its predictive performance, we asked whether SubNetDL could be used to identify genes that contribute to immunotherapy response—i.e., candidate biomarkers distinguishing responders from non-responders. Building on our earlier analysis in the TCGA-derived datasets, we applied the same gene importance framework to the MSK_2018 and MSK_2020 cohorts. For each dataset, we selected the top 5% of genes based on importance scores (12 genes for MSK_2018; 12 genes for MSK_2020), which retained 0.95 ratio of the original model’s performance (top 2.5% = 0.84; top 10% = 1 ratio of the original model’s performance). Notably, the top 5% genes were repeatedly identified across cross-validation folds and in both MSK_2018 and MSK_2020 cohorts, supporting their robustness and potential as reproducible biomarkers (**Fig. 4E**). Several of the top-ranked genes identified by SubNetDL, such as *ARID5B*, *ZFP36L2*, and *POLQ*, have previously been linked to immunotherapy response through their roles in cytokine regulation, immune cell development, and DNA damage signaling^47–51^. In addition, *RNF43* and KDM5C have been associated with immune checkpoint blockade outcomes in MSI-high tumors and NSCLC, respectively. Notably, *CNBD1*, which ranked first in both cohorts, has not been directly implicated in immunotherapy response. However, it was recently reported as part of an immune cell–associated gene signature in gastric cancer, suggesting a potentially unexplored role in modulating tumor–immune interactions^52^. These findings suggest that SubNetDL not only enhances prediction accuracy but also provides biologically meaningful insights, enabling the identification of treatment-specific gene signatures with potential translational value.

To further evaluate whether SubNetDL generalizes across distinct biological systems and response readouts, we performed an independent validation using the Cancer Cell Line Encyclopedia (CCLE)^53^, with drug response measurements from the Genomics of Drug Sensitivity in Cancer (GDSC) resource. Unlike patient-derived TCGA and MSK cohorts, CCLE represents a preclinical model system with experimentally measured IC50 values. Applying the same modeling framework without cohort-specific tuning, SubNetDL achieved strong predictive performance across six cancer–drug pairs (median AUROC = 0.81, range: 0.70–0.85; **Supplementary Figure 4C and D**).

## Discussion

While therapeutic response prediction has been a growing focus over the past 15 years—particularly following the emergence of immune checkpoint inhibitors—many existing approaches have focused on predefined biomarkers or mutation burden alone^5^. Here, we present a network propagation framework that integrates functional relationships among genes to infer subclonal architecture. By addressing subclonality at the patient level, our approach captures the dynamic and context-dependent nature of tumor evolution, including clonal shifts under therapeutic pressure. These clonal features are not only predictive of drug response but may also inform broader clinical implications, such as treatment resistance and disease progression^23,24,54^. Unlike condition-specific models that rely on curated gene sets, our method requires only patient mutation profiles and a generic human protein interaction network, allowing for broader applicability across cancer types and treatment modalities.

SubNetDL distinguishes itself from conventional network-based approaches that prioritize genes based on static topological features—such as centrality or local connectivity. To evaluate this difference, we compared SubNetDL’s gene-level importance scores against six classical topology metrics, including degree, betweenness, closeness, eigenvector centrality, clustering coefficient, and average neighbor degree. In all comparisons, correlations were weak or inconsistent (Person correlation coefficient ranging from -0.15 to 0.21), suggesting that SubNetDL does not simply rely on hub proteins or topologically central nodes (**Fig. 5A**). Specifically, we directly compared SubNetDL’s gene importance rankings with network degree (i.e., the number of interacting partners), where a top-ranked gene corresponds to a hub protein in the PPI network. Interestingly, we observed an inverse association between SubNetDL scores and degree-based rankings (**Fig. 5B** and **Supplementary Figure 4E**). Notably, canonical hub genes—such as *TP53*, *AKT1*, and *EGFR*—were not ranked within the top 5% by SubNetDL and were not prioritized highly. This underscores that SubNetDL captures higher-order, context-specific patterns beyond conventional network centrality.

**Figure 5.**
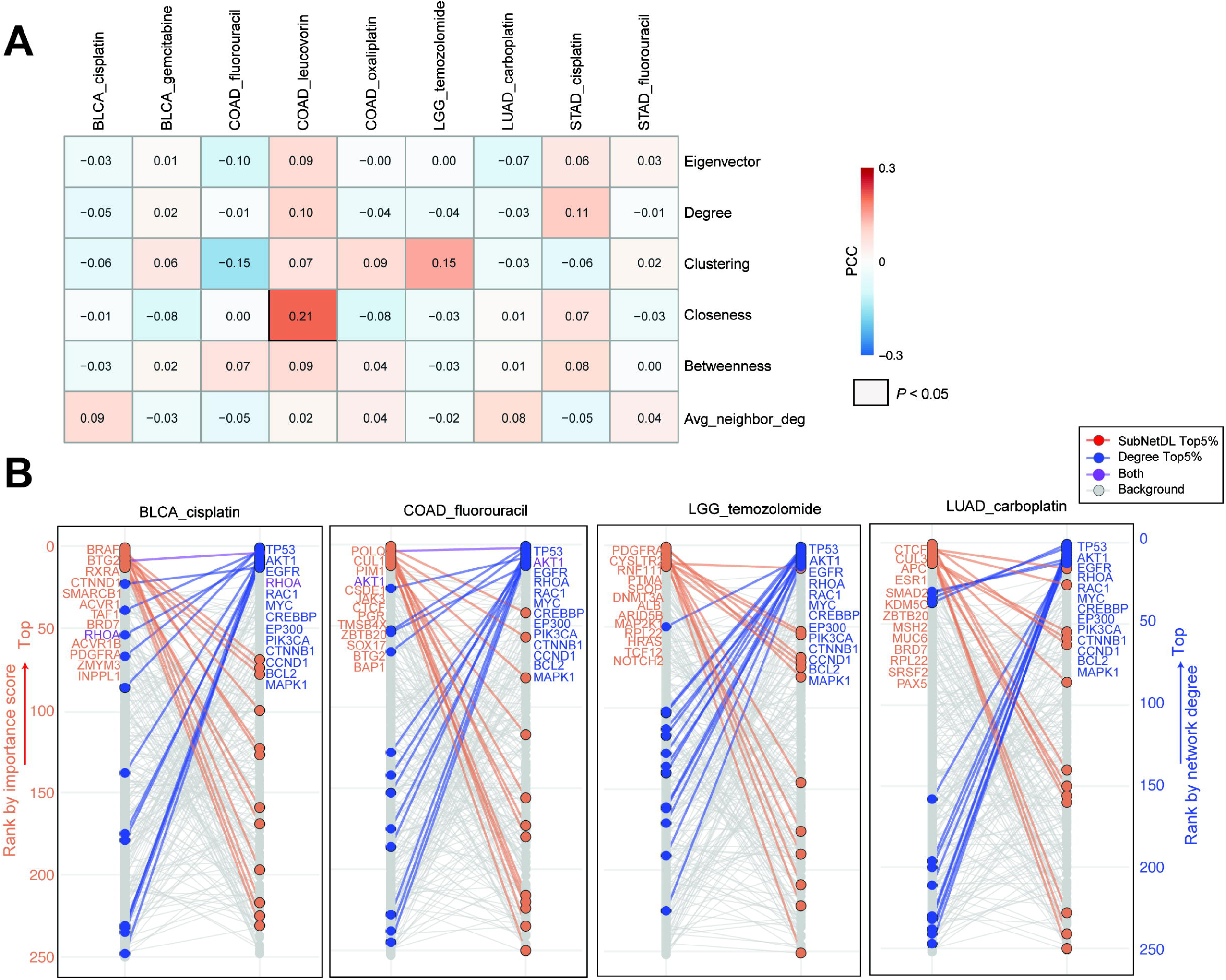
Comparison of gene ranks based on SubNetDL importance scores and network degree across cancer type–treatment pairs. (**A**) Overview of person correlation coefficient between importance score from SubNetDL and six network-driven features. (**B**) Genes ranked in the top 5% by SubNetDL are labeled in red, those ranked in the top 5% by network degree (based on the STRING PPI network) are labeled in blue, and genes ranked in the top 5% by both are shown in purple. All other genes are shown in grey. Four cancer type–drug combinations are presented here; at least five additional combinations are provided in **Supplementary Figure 4E**.

Beyond performance gains, SubNetDL offers a framework for identifying mechanistically relevant features. For instance, *CNBD1* consistently ranked as the top predictive gene across cohorts, despite limited prior links to immunotherapy. A recent study reported its inclusion in an 11-gene signature associated with immune infiltration and gastric cancer prognosis^52^, pointing to a potentially underexplored role in tumor–immune interactions. Similarly, other recurrently prioritized genes—such as *ARID5B*, *ZFP36L2*, *RNF43*, *POLQ*, and *KDM5C*—have been implicated in cytokine regulation or immune checkpoint activity^47–51^, suggesting that SubNetDL may enrich for treatment-relevant biology.

Although we did not observe a clear correlation between the average number of subclones per cancer type and predictive performance (*r* = -0.17 and *P* = 0.64; **Supplementary Figure 5A**), subclonal architecture remains a critical component of our method. To assess the model’s dependence on subclonal reconstruction, we performed perturbation analyses in which mutations were randomly reassigned across inferred subclones within each patient while preserving overall mutation burden. Disrupting clonal organization significantly reduced predictive performance, indicating that SubNetDL leverages biologically coherent mutation groupings rather than relying solely on aggregate mutational features (median AUROC = 0.73 vs. 0.58, *P* = 0.02; **Supplementary Figure 5B** and **C**). In contrast, restricting the model input to only the dominant subclone per patient did not introduce systematic prediction bias, suggesting that the framework is not overly sensitive to low-prevalence clonal fluctuations under bulk sequencing resolution (median AUROC = 0.74, *P* = 0.43; **Supplementary Figure 5B** and **C**).

To further contextualize subclonal heterogeneity, we examined its relationship with established molecular subtypes in STAD and COAD, where subtype annotations are well defined. Subclonal complexity varied across subtypes, with genomically stable contexts showing lower subclone counts compared to chromosomally unstable or hypermutated groups. Treatment response distributions likewise differed by subtype, and predictive performance tended to be more stable in molecularly homogeneous contexts. These observations suggest that subtype stratification provides important context for interpreting both subclonal architecture and response prediction (**Supplementary Figure 5D-F**).

To ensure the reliability of these findings, we confirmed that model performance remained stable under moderate perturbations of key design parameters, including sequencing depth, propagation scale, network depth, and driver gene set size (**Supplementary Figure 5G-M**). Alternative configurations did not materially alter predictive accuracy, indicating that SubNetDL is not driven by narrowly tuned hyperparameters. This robustness supports that the predictive signal arises from biologically meaningful structure rather than flexible model capacity. In conclusion, SubNetDL does not merely reproduce known predictors but learns functionally meaningful patterns that generalize across therapeutic contexts. By operating on widely available mutation data without requiring condition-specific training, SubNetDL offers a flexible and interpretable alternative to classical gene-centric models—one that can uncover novel biomarkers while enhancing prediction accuracy.

## Resource Availability

### Lead Contact

Requests for further information and resources should be directed to and will be fulfilled by the lead contact, Solip Park.

### Materials Availability

The study did not generate new unique reagents.

### Data and Code Availability

All data used in this study are publicly available. TCGA was downloaded by R package

TCGAbiolinks. Data for two immunotherapy treated cohort was downloaded from cBioPortal. The human PPI network was downloaded from STRING (https://string-db.org/). All data needed to evaluate the conclusions in the paper are present in the paper and/or the Supplementary Materials.

- Code: All original code has been deposited at http://github.com/SolipParkLab/SubNetDL (https://zenodo.org/records/18982291).
- Any additional information required to reanalyze the data reported in this work is available from the lead contact upon request.

## Limitations of the study

Our current approach for estimating subclonality has important limitations. First, subclonality is inferred from bulk sequencing data, which limits the resolution of clonal structures—especially in the presence of spatial or temporal heterogeneity^23,24^. While bulk-based inference is supported by prior studies^32,55,56^, it cannot fully resolve fine-grained clonal compositions. Recent single-cell approaches offer higher resolution and are increasingly used to uncover complex tumor evolution dynamics^57–59^, although variant calling from single-cell data remains technically challenging^60–62^. We anticipate that integrating high-resolution subclonality profiles—through single-cell or spatial data—will further enhance the performance and interpretability of SubNetDL. Second, our subclonality estimation is primarily driven by mutation count distributions, which may be difficult to detect clones present at low frequencies or those subject to weak selection. Such clones may be transient, spatially restricted, or below the detection limit of standard sequencing depths. In such settings, the inferred clonal architecture may not fully capture rare or weakly selected populations. Future efforts that incorporate longitudinal sampling or cell-free DNA profiling may help overcome these limitations. Finally, while we demonstrated the robustness of SubNetDL across multiple cohorts, the current framework relies on a generic human protein interaction network. While this allows for broader applicability, incorporating tissue-specific or context-dependent networks could potentially refine the functional relationships identified by the model.

## Acknowledgements

The results shown here are in whole or part based upon data generated by the TCGA. A.N. is supported by the AI-Cancer funded from the European Union’s Recovery and Resilience Facility-Next Generation, in the framework of the General Invitation of the Spanish Government’s public business entity Red.es to participate in the talent attraction and retention programmes within Investment 4 of Component 19 of the Recovery, Transformation and Resilience Plan. S.K. is supported by grant from the Korean National Research Foundation (2020R1A6A1A03047902). S.P. is supported by the Korean National Research Foundation of (RS-2024-00415982) grant. S.P. is supported by the project PID2019-109571RA-I00 and PID2022-141202OB-I00 funded by the Agencia Estatal de Investigación (AEI/10.13039/501100011033), Ministerio de Ciencia e Innovación and co-founded by the European Regional Development Fund (ERDF-EU); and CNS2023-144982 funded by MCIN/AEI/10.13039/501100011033 and co-founded by the European “NextGenerationEU”/PRTR. S.P. is also supported by Ramón y Cajal grant RYC2021-034415-I, funded by MCIN/AEI/10.13039/501100011033 and co-founded by the European Union – “NextGenerationEU”/PRTR and Ramón Areces Foundation. S.P. acknowledges the support of the Severo Ochoa Centres of Excellence program to the CNIO CEX2019-000891-S funded by the MCIN/AEI/10.13039/50110001103.

## Author Contributions

Conceptualization, S.N.K., J.H.L., S.K., and S.P; methodology, S.N.K., J.H.L., S.K., and S.P.; software, S.N.K., D.H., J.H.L.; validation, S.N.K.; formal analysis, S.N.K., A.N., J.H.L., S.K., and S.P.; investigation, S.N.K., J.H.L., S.K., and S.P.; resources, S.N.K., D.H., S.C., J.H.L.; data curation, S.N.K., D.H., S.C., J.H.L..; writing–original draft, S.N.K., J.H.L., S.K., and S.P.; writing–review & editing, S.N.K., J.H.L., S.K., and S.P.; visualization, S.N.K., A.N., and S.P.; supervision, J.H.L., S.K., and S.P.; funding acquisition, S.K., and S.P.

## Declaration of Interests

J.H.L. is employee of ImmunoBiome. The remaining authors declare no competing interests.

## STAR Methods text

### EXPERIMENTAL MODEL AND STUDY PARTICIPANT DETAILS

#### TCGA-based genomic and clinical data collection

In this study, we utilized publicly available multi-omics data from The Cancer Genome Atlas (TCGA), comprising a cohort of 11,167 cancer patients. Three types of data—clinical information, Copy Number Variation (CNV), and Single Nucleotide Variant (SNV)—were retrieved using the R package TCGAbiolinks (version 2.16.4)^63^. CNV data were downloaded via the GDCquery function using the following parameters: project = ‘cancer_type’, data.category = ‘Copy Number Variation’, data.type = ‘Copy Number Segment’, and sample.type = ‘Primary Tumor’. The resulting dataset provided copy number alterations for specific chromosomal regions represented as Segment Mean values, encompassing 9,797 samples. SNV data were obtained using the GDCquery_Maf function with pipelines = ‘muse’ (10,489 samples). The MuSE pipeline (Mutation Calling Using a Markov Substitution Model for Evolution) accounts for tumor heterogeneity by employing a sample-specific error model^64^. Variants were annotated by MuSE into categories such as missense_mutation, silent, 5’Flank, 3’UTR, RNA, nonsense_mutation, splice_site, intron, 5’UTR, nonstop_mutation, 3’Flank, and translation_start_site. For each variant, MuSE also provides the number of reads supporting the reference allele (REF count) and the alternative allele (ALT count). Clinical information was retrieved using the GDCquery function with the following arguments: project = ‘cancer_type’, data.category = ‘Clinical’, data.type = ‘Clinical Supplement’, and data.format = ‘BCR Biotab’. The column ‘pharmaceutical_therapy_drug_name’ was used to extract drug names from the treatment history provided in the dataset (available for 4,328 samples). To classify drug response, we used the column treatment_best_response, which indicates the best observed clinical outcome for each treatment (available for 1,701 samples). Based on this, patients were categorized into responders (Complete Response [CR] and Partial Response [PR]) and non-responders (Stable Disease [SD] and Progressive Disease [PD]). After integrating clinical response data with matched SNV and CNV profiles, a total of 1,470 patients were included in the final analysis (**Supplementary Table 1**). In cases where patients received multiple therapies, only the final treatment and its associated response were retained for sequential regimens (*n* = 7), whereas combination therapies were decomposed into independent patient–drug instances (*n* = 8), with all train–test splits performed at the patient level to prevent data leakage (**Supplementary Table 2**).

Clinical survival data, including overall survival (OS) and progression-free interval (PFI), were obtained from the TCGA Pan-Cancer Clinical Data Resource^65^ through the GDC PanCanAtlas portal.

## METHOD DETAILS

### Subclonal structure estimation

To infer subclonal architectures in individual cancer patients, we employed the R package SciClone (version 1.1.1; R version 3.6.3; https://github.com/genome/sciclone) with default parameters. The input for SciClone included the variant allele frequency (VAF) of functional exonic SNVs—calculated as ALT count / (REF count + ALT count)—alongside copy number segment means for each genomic region. SciClone utilizes a variational Bayesian beta mixture model to cluster SNVs based on their posterior predictive density, thereby grouping SNVs with similar VAF values into subclones. To correct for regional differences in copy number, the expected VAF for each SNV is adjusted according to the segment mean in the corresponding copy-number region. Adjusted VAFs are used for clustering, but variants that do not meet filtering criteria are excluded from subclonal cluster assignment. To ensure reliability, SciClone excludes clusters containing fewer than 3 SNVs or accounting for less than 0.5% of all variants, depending on which cutoff is more restrictive. For example, with 1,000 SNVs, any cluster with fewer than 5 variants (0.5%) would be excluded before re-clustering. This iterative process continues until statistically robust subclonal clusters are identified. SciClone has been benchmarked to identify stable and biologically relevant subclonal clusters typically within the range of 1 to 6, which we adopted for downstream analyses. Subclonal clustering may not be achieved when the input data lacks sufficient signal—such as limited SNV counts, low read depth, or tightly clustered VAF distributions—conditions under which distinct subclones cannot be resolved. Among the 1,470 patients with genomic data, subclonal architecture could be inferred in 944 cases (**Supplementary Table 1**).

### Development of a SubNetDL model

We developed a network-based deep learning model, aiming to therapeutic response prediction in cancer patients by incorporating the functional association strengths among subclonal somatic variants. The model is based on Graph Attention Networks (GATs)^34^.

#### - Input: network-propagated scores seeded from SNVs in a subclone

Human protein-protein interaction (PPI) network from the STRING database (version 11.0; https://string-db.org/)^33^, collecting only high-confidence interactions with a combined score greater than 700, as described by Fernández-Torras *et al*.^66^, was used for the backbone network following analysis. This resulted in a network of 255,825 interactions among 14,278 human proteins. For each subclone, we mapped binary SNV information onto the network as initial values (1 for proteins with an SNV, 0 otherwise). We then ran propagation analyses, and the propagation score of a gene was utilized as functional association strength of mutated genes in the subclone. Furthermore, we used five different damping factors (0.05, 0.25, 0.45, 0.65, and 0.85) to control the extent of the association strength across the network. A lower damping factor limits propagation to the immediate neighborhood, emphasizing local mutation clustering. In contrast, higher values allow propagation to more distant nodes, thereby reflecting the network-central position of a mutated protein. By applying a range of damping factors, we aimed to capture both local SNV density and global topological centrality (**Supplementary Figure 1A**). Network propagation was performed using the PageRank algorithm^67^, implemented via the NetworkX Python package (Python 3.7.10, NetworkX 2.5). Originally developed to rank web pages based on their link structure, PageRank has been successfully applied in biological contexts to identify novel biomarkers and predict therapeutic response in cancer patients^30^.

#### - Model architecture

The proposed model consists of two main modules: an embedding module and a classification module. Both modules are implemented using GATs^34^, built upon the STRING functional PPI network. The embedding module consists of two GAT layers, while the classification module includes one GAT layer followed by a pooling layer and a fully connected layer. GAT layers enable learning from interactions among genes (nodes) by assigning attention weights between connected proteins, allowing phenotype prediction based on network-aware mutation context. The mathematical definition of a GAT layer is as follows:

For intermediate layers:

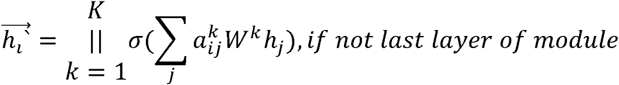

For the last layer of each module (averaged heads):

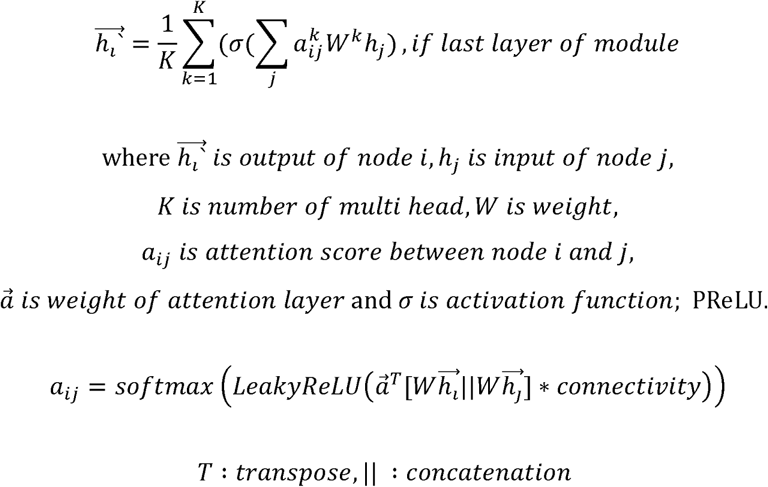

The multi-head attention mechanism was implemented as described by Veličković *et al*.^34^. In the embedding module, the first GAT layer used 5 heads (concatenated outputs), and the second layer used 3 heads (averaged outputs). The classification module used a single-head GAT layer to facilitate interpretability.

To focus on cancer-relevant biology and reduce the risk of overfitting due to excessive model complexity, we pruned the STRING network to include only 253 known cancer driver genes and used it for training. To preserve the global connectivity of the network and account for the topological proximity between genes, we redefined the weight of each link as the inverse of the shortest path length between gene pairs (i.e., connectivity = 1 / shortest path distance).

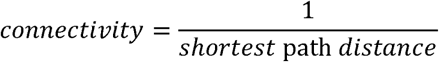

These 253 genes, overlapping with nodes in the STRING network, were selected from a list of 299 cancer driver genes curated by Bailey *et al*.^35^ based on 9,423 tumor exomes analyzed with 26 computational tools. To incorporate node proximity, we assigned edge weights as the inverse of the shortest path distance between two nodes. This weighting encourages greater information exchange between nearby nodes in the graph^34^. For each of the 253 genes, five propagation scores were generated per subclone. Across all subclones in a patient (ranging from 1 to 6), the input to the model was structured as a 3D tensor:

[number of subclones] × [253 genes] × [5 damping factors].

This formulation allows nodes that are more closely connected to exchange more information during message passing. Layer specifications were as follows:

Embedding module (per subclone and per gene):

Layer 1: input size = 5 (feature vector size per gene), output = 10 (×5 heads = 50) Layer 2: input = 50, output = 10

Classification module:

GAT layer (per subclone and per gene): input = 10, output = 1

Pooling layer: input = 253 × number of subclones, output = 253

Final fully connected layer: input = 253 (pooled from all subclones), output = 1 (binary prediction)

Each patient sample typically contains between one and six subclones, making the model design non-trivial due to the variable number of subclonal inputs. To accommodate this variability, SubNetDL applies a parameter-shared Graph Attention Network independently to each inferred subclone. This ensures consistent feature extraction regardless of subclone number. The embedding module maps each subclone to a fixed-length vector that summarizes its mutational profile. The outputs (one embedding per subclone) are then aggregated into a single patient-level representation using a pooling layer in the classification module. This pooled vector is subsequently used to predict drug response (responder versus non-responder).

#### - Datasets for training and evaluating of a subclonal architecture-aware deep learning model

To train a deep learning model for predicting patient drug response (i.e., responder *versus* non-responder), we grouped patients according to their cancer type and received therapy. To ensure model stability, we considered only cancer type–drug pairs with at least 30 patients in total. Among these, we selected the top 10 cancer type–drug groups with the largest number of patients in the minority class—defined as the smaller of the responder or non-responder groups (**Supplementary Table 3**). A total of 506 patients with known drug response status (responder or non-responder) were included for supervised learning. These patients spanned six cancer types (BLCA, COAD, STAD, LUAD, LGG, CESC) and received one of seven drugs: cisplatin, fluorouracil, gemcitabine, leucovorin, oxaliplatin, carboplatin, or temozolomide. Sample sizes per group ranged from 30 to 71.

### Training and evaluation of the network-based deep learning model

For each cancer–drug pair, patients were randomly split into training (80%), validation (10%), and test (10%) sets using a fixed random seed (42) for reproducibility. The model was trained using stochastic gradient descent with the Adam optimizer (learning rate = 1e-6, batch size = 1) and employed the PReLU activation function^68,69^. Dropout (rate = 0.2) was applied after each fully connected layer to reduce overfitting. All weights were initialized using Xavier normal initialization to maintain activation variance across layers and promote stable convergence. As the model was designed to predict binary response outcomes (responder *versus* non-responder), Binary Cross-Entropy Loss was used as the objective function. The final model was selected based on the highest AUROC achieved on the validation set and evaluated on the test set to report predictive performance.

### Ablation analysis of subclone inference and network-based deep learning for predictive modeling

To assess the individual contributions of subclone inference and the network-based deep learning framework, we conducted ablation experiments by removing each component from the model. In the model without subclone inference, functional exonic SNVs from each patient were not divided into subclones. Instead, all SNVs were mapped onto a single network using binary indicators (1 for presence, 0 for absence), and this network representation was used as input to the deep learning model. In the model without network-based deep learning, we retained the subclone assignments but replaced the deep learning framework with conventional machine learning classifiers, including support vector machines (SVM), random forest, linear regression, and logistic regression. The random forest model was configured with n_estimators = 1,000, max_depth = 10, and random_state = 0. The SVM was applied with a linear kernel. All other models were used with default hyperparameter settings.

### Attention-based gene prioritization for anti-cancer therapy response prediction

To identify genes contributing to the prediction of drug response, we analyzed the attention weights from the GAT in the classification module. In GAT, the attention coefficient a*_ij_* between gene *<u>i</u>* and gene *j* represents the importance of node *j* ‘s feature to node *i*—that is, how much the feature of gene *j* influences the updated representation of gene *i*. Since the final GAT layer aggregates the learned representations from all previous layers and directly informs the model’s output, we focused on this layer for interpretation. For each patient, we extracted all pairwise attention values a*_ij_* from the final layer. A larger a*_ij_* indicates that the feature of gene *j* has a stronger influence on gene *i*’s representation and is therefore more important for prediction. We define the importance of gene *j* as follows:

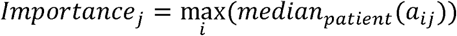

We first identified gene pairs (*i, J*) whose attention weights significantly differed between responders and non-responders (Bonferroni-adjusted *P* < 0.01). For each such gene *j*, we computed the median a*_ij_* across all patients and took the maximum across all *i* to quantify its importance. Genes were then ranked by their importance scores. We trained new deep learning models using only the top-ranked genes (e.g., top 5%) as input and found that these models achieved performance comparable to the full model across most cancer–drug combinations (**Fig. 3**). Finally, we performed functional enrichment analysis on the top 5% of genes using DAVID^70^, collecting enriched Gene Ontology Biological Process (GO:BP) and Molecular Function (MF) terms with adjusted *P*-values below 0.01.

To compare genes identified by attention weights with those identified through network topological analysis, we computed node centrality and structural metrics using the NetworkX Python package (version 3.1). Based on the STRING (with integrated score over 700) PPI network used for network propagation, we calculated degree centrality, betweenness centrality, closeness centrality, eigenvector centrality, clustering coefficient, and average neighbor degree. For the calculation of eigenvector centrality, the max_iter parameter was set to 1,000, with all other parameters kept at their default values.

### Evaluating predictive performances using gene expression-based biomarkers

Gene expression data for cancer patients were downloaded from the TCGA using the GDCquery function with the following parameters: project = ‘cancer_type’, data.category = ‘Transcriptome Profiling’, workflow.type = ‘HTSeq - Counts’, and sample.type = ‘Primary Tumor’. A total of 507 TCGA patients were included for gene expression marker-based performance analysis. To benchmark our model performance, we curated known expression-based drug response biomarkers from the literature searches (**Supplementary Table 4**). AUROC was calculated for each biomarker using patient gene expression levels to classify responders versus non-responders.

### Evaluation of a subclonality-aware deep learning model in an immunotherapy-treated cohort

We downloaded genomic and drug response data for 247 and 240 non-small cell lung cancer (NSCLC) patients treated with PD-(L)1 blockade therapy from Vanguri., *et al*.^46^ and Rizvi *et al*.^45^, respectively. In both studies, tumor and matched normal specimens were analyzed using the MSK-IMPACT assay, a clinically validated hybrid capture-based sequencing panel covering 341–468 cancer-associated genes. All patients received PD-(L)1 blockade therapy. Mutation, CNV, and response data were obtained from cBioPortal(https://www.cbioportal.org/study/summary?id=lung_msk_mind_2020, https://www.cbioportal.org/study/clinicalData?id=nsclc_pd1_msk_2018). For mutations, we used all coding region mutations provided in the data_mutations.txt file. For CNVs, we used the seg.mean values from the data_cna_hg19.seg file. Clinical information was obtained from the data_clinical_patient.txt file. In the Vanguri *et al*. cohort, we defined 61 patients with a BOR (Best Overall Response) status of CR (Complete Response) or PR (Partial Response) as responders, and the remaining 186 patients as non-responders. In the Rizvi *et al*. cohort^46^, we defined 42 patients with a “Progression Free Status” of “0: Not Progressed” as responders and 198 patients with “1: Progressed” as non-responders.

## QUANTIFICATION AND STATISTICAL ANALYSIS

All statistical analyses were performed using Python (version 3.7.10). Genomic and clinical data were retrieved using the TCGAbiolinks R package, while network topological metrics and PageRank propagation were implemented via the NetworkX Python package. Patient-specific subclonal architectures were inferred using SciClone (version 1.1.1; R version 3.6.3). For supervised learning, model stability was ensured by including only cancer type–drug pairs with at least 30 samples, resulting in 506 patients across ten TCGA pairs and external validation in MSK_2020 (n = 242) and MSK_2018 (n = 240) immunotherapy cohorts. Data were randomly partitioned following an 8:1:1 stratified split for training, validation, and testing of TCGA cohorts to preserve class balance, while 10-fold cross-validation was implemented for independent NSCLC cohorts to ensure robust assessment. Model performance was primarily evaluated using the area under the receiver operating characteristic curve (AUROC), supplemented by accuracy, the area under the precision-recall curve (AUPRC), and F1 score. Predictive performance differences between SubNetDL and ablation settings or classical machine learning models were assessed using paired two-sided Wilcoxon signed-rank tests. Clinical survival differences based on prediction scores were analyzed using Kaplan–Meier survival curves and log-rank tests. Functional enrichment for significant genes prioritized by the model (top 5%) was determined via Fisher’s exact test for Gene Ontology (GO) terms, and the relationship between attention-based gene importance scores and network topology metrics was quantified using Pearson correlation coefficients. Statistical significance was defined as P < 0.05 across all tests, and data summaries in figures were presented as medians, ranges, or interquartile ranges (IQR) where applicable.

## supplemental video and Excel table titles and legends

**Supplementary Table S1**. Summary of TCGA patient cohorts and sample inclusion criteria across sequential filtering steps, related to STAR Methods. This table details the sample size transitions for 33 TCGA cancer types during the data processing pipeline. It tracks the number of patients from initial data collection through the availability of matched SNV/CNV profiles, successful subclonal architecture inference using SciClone, and the presence of clinical treatment and response records.

**Supplementary Table S2**. Identification and processing of patients receiving combination therapies, related to STAR Methods. This table lists specific patient instances from the COAD and BLCA cohorts who received combination therapy regimens (e.g., fluorouracil, leucovorin, and oxaliplatin). It describes how these cases were decomposed into independent patient–drug instances for the SubNetDL framework.

**Supplementary Table S3**. Sample size distribution and cohort selection for therapeutic response analysis, related to Figure 2. Detailed distribution of responders and non-responders across various TCGA cancer–drug combinations. This table highlights the selection process for the ten primary evaluation cohorts, which were chosen based on a minimum sample size of 30 and the largest representation of the minority class (the smaller of the responder or non-responder groups).

**Supplementary Table S4**. Literature-curated gene expression-based biomarkers for anti-cancer drug response, related to Figure 2. A comprehensive list of established expression-based biomarkers and signatures curated from clinical literature. These biomarkers were utilized to benchmark the predictive performance of SubNetDL’s mutation-informed framework against traditional gene expression-based classifiers.

## Notes

https://github.com/SolipParkLab/SubNetDL

