## Supplementary Figures for "A network-based deep learning model integrating subclonal architecture for therapy response prediction in cancer"

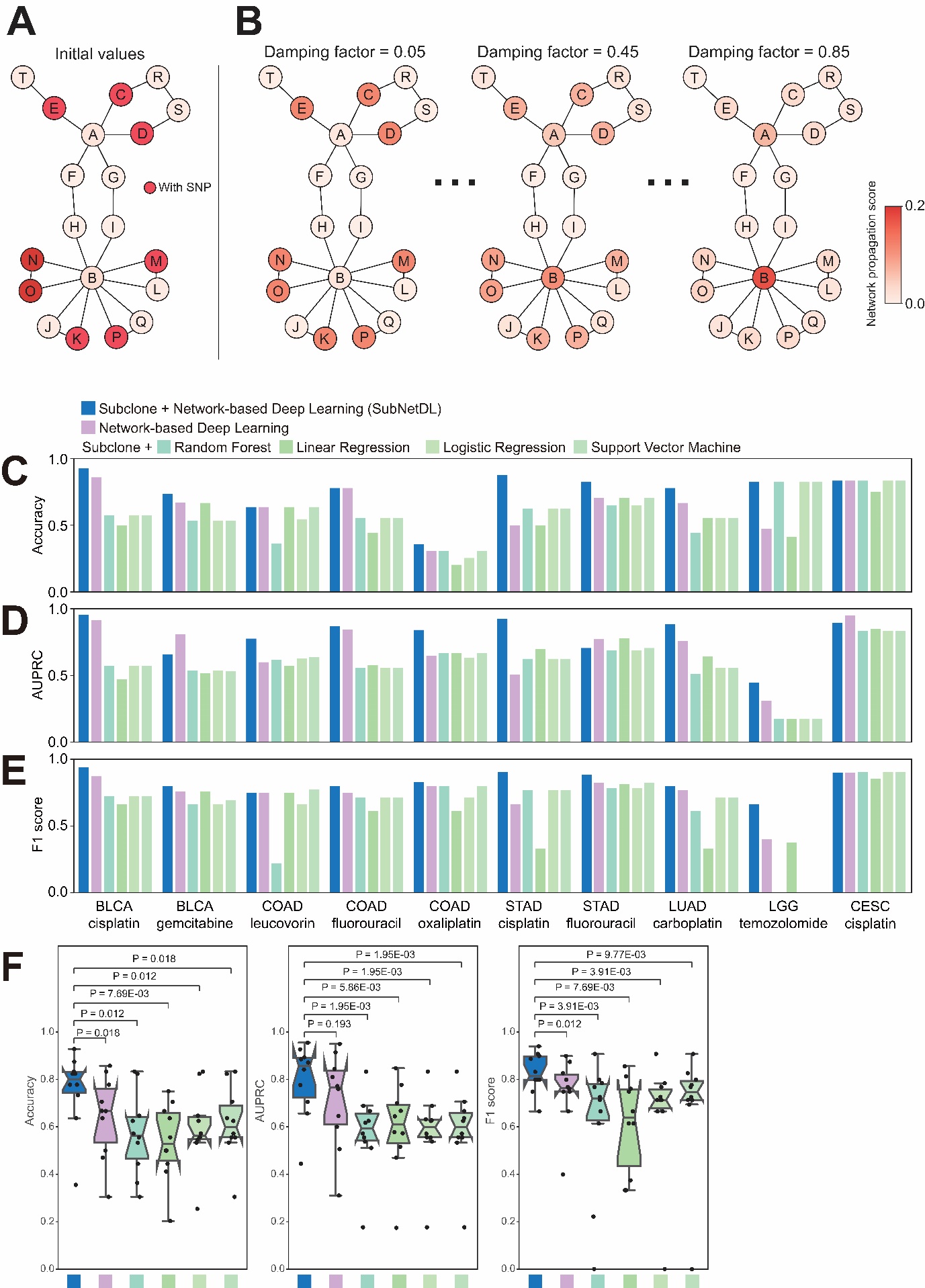

**Supplementary Figure 1**. **SubNetDL leverages multiscale mutation propagation to predict cancer drug response, related to Figure 1. (A) Binary mutation labels were assigned to nodes in a protein–protein interaction network (1 = mutated, red; 0 = not mutated, white), with red nodes indicating the presence of a mutation. (B) Mutation-informed network propagation was applied using three example damping factors (α = 0.05, 0.45, 0.85), enabling the diffusion of mutational signals from initially mutated nodes to their neighbors. Lower α values (e.g., 0.05) emphasize local interactions, while higher values (e.g., 0.85) allow broader signal spread. Node color intensity reflects the resulting propagation scores, with darker red indicating stronger influence from mutated nodes.** (**C**–**E**) Performance comparison of six models across three evaluation metrics: (**C**) Accuracy, (**D**) AUPRC, and (**E**) F1 score, using an 8:1:1 split for training, validation, and testing. (**F**) Accuracy, AUPRC, F1 score distributions across the ten cancer–drug pairs for each model. Each point represents a single cancer–drug combination. Box plots display the interquartile range (IQR), with center lines indicating the median and whiskers extending to 1.5×IQR. Statistical comparisons were performed using paired two-sided Wilcoxon signed-rank tests.

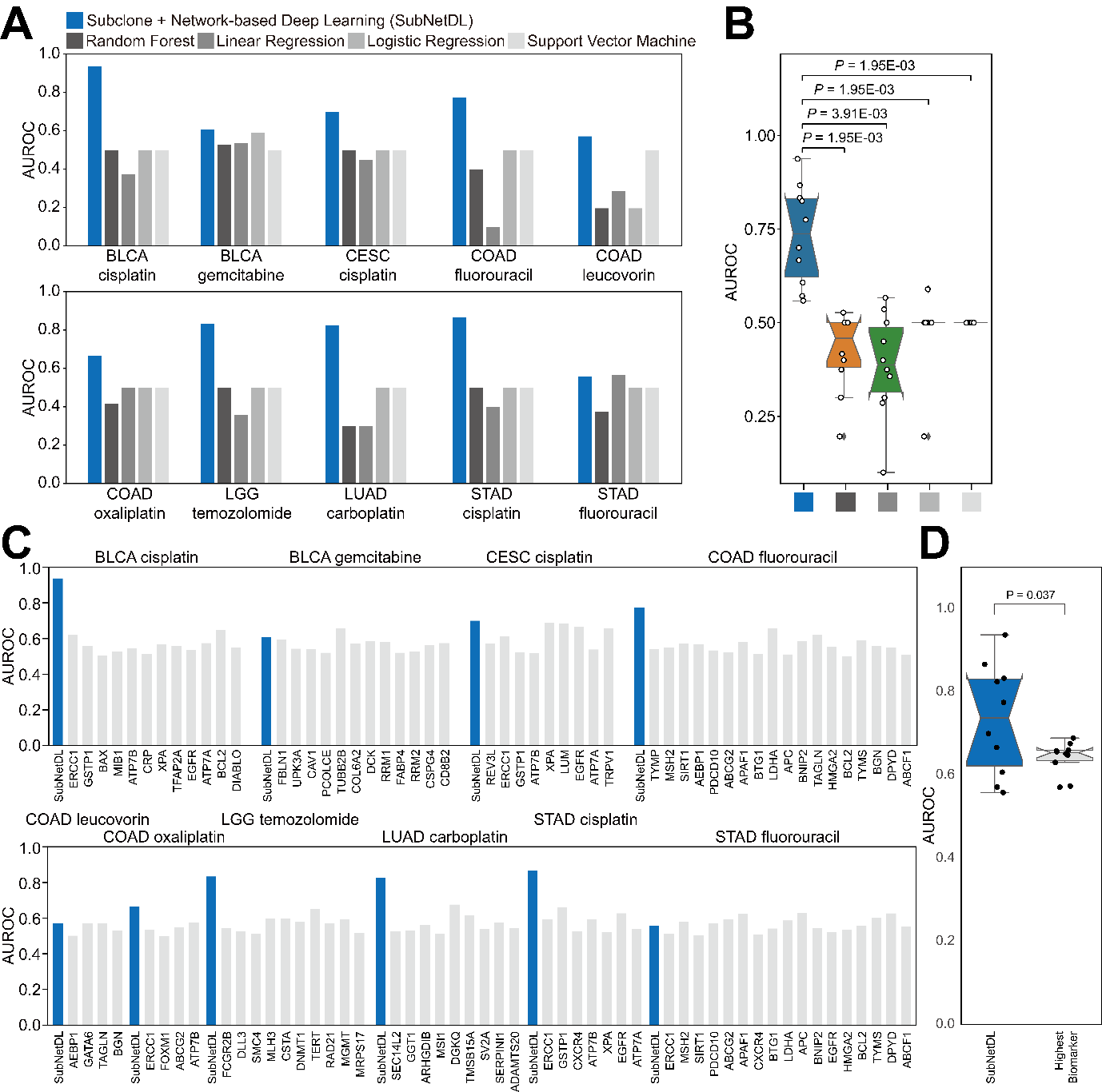

**Supplementary Figure 2**. Comparative predictive performance of SubNetDL across ten cancer–drug treatment pairs**, related to Figure 2**. (**A**) Performance (AUROC) comparison of five models. (i) subclonality-integrated network-based deep learning model (blue; SubNetDL), (iii) four mutation-based machine learning models using random forest, linear regression, logistic regression, and support vector machine (gray, without subclone and graph-based deep learning). Models are evaluated using the same 8:1:1 train–validation–test split. (**B**) Distribution of AUROC scores across the ten cancer–drug pairs for each model. Each point represents a single cancer–drug pair. Box plots indicate the interquartile range (IQR), with center lines denoting the median and whiskers extending to 1.5×IQR. Statistical comparisons were performed using paired two-sided Wilcoxon signed-rank tests. (**C**) Comparison of SubNetDL performance with expression-based biomarkers previously reported to predict drug response in cancer patients, using AUROC as the evaluation metric. (**D**) Boxplot summarizing AUROC scores between the ten cancer–drug pairs and expression-based biomarkers with the highest AUROC across cancer types. Boxes represent the interquartile range (IQR), center lines denote the median, and whiskers extend to 1.5×IQR. Statistical comparisons between models were performed using paired two-sided Wilcoxon signed-rank tests. *P*-values < 0.05 and < 0.005 are indicated by * and **, respectively.

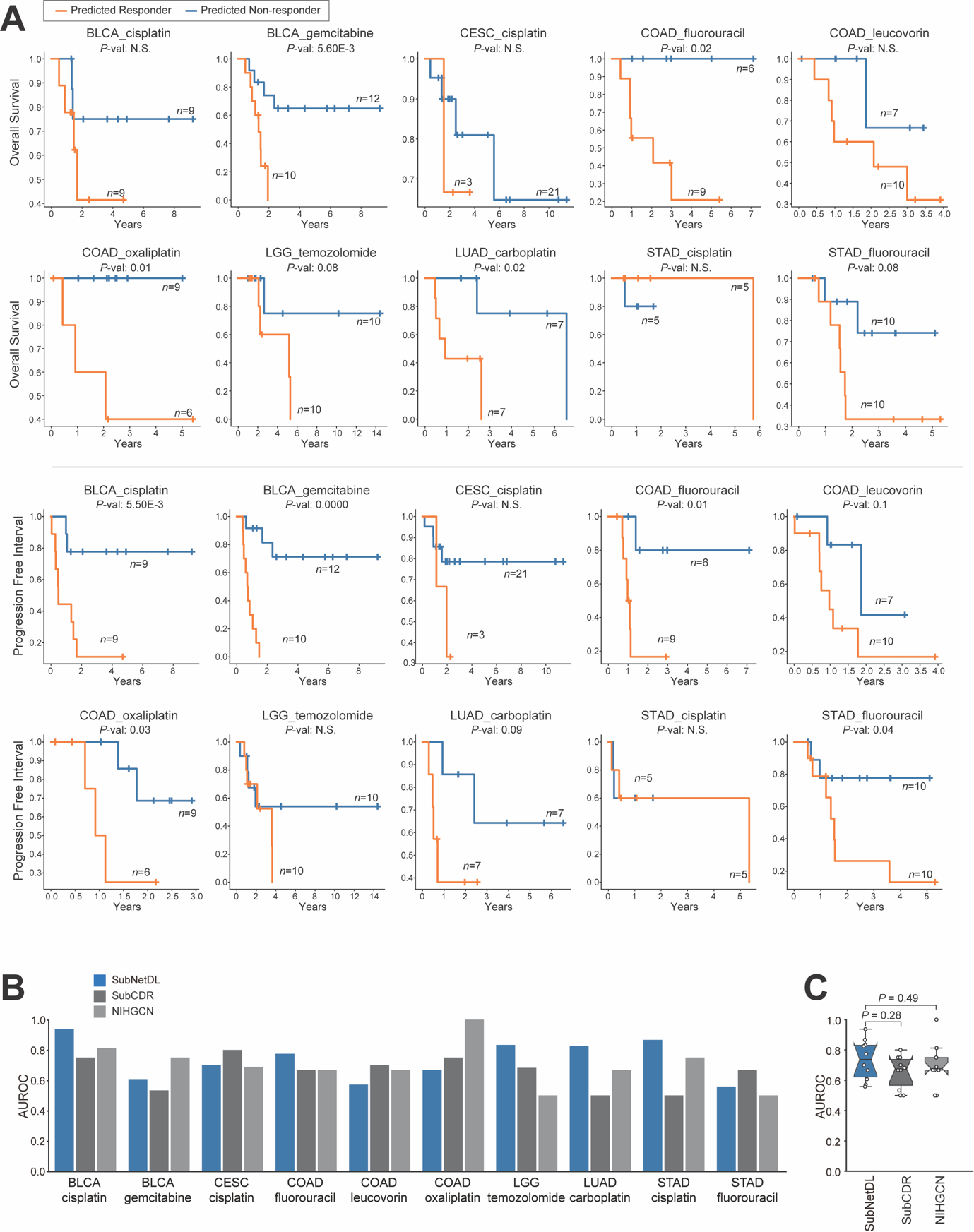

**Supplementary Figure 3**. SubNetDL predictions associate with patient survival and improve drug response prediction**, related to Figure 2**. (**A**) Kaplan–Meier curves show Overall Survival (OS; top two rows) and Progression-Free Interval (PFI; bottom two rows) across ten TCGA-derived cancer–drug combinations. Patients were stratified according to the median SubNetDL prediction score into high-score and low-score groups. For each cohort, the number of patients (*n*) per group and the corresponding log-rank test *P*-value are indicated. (**B**) AUROC comparison of three predictive frameworks: SubNetDL (blue), a subclonality-integrated network-based deep learning model; SubCDR (dark gray); and NIHGCN (light gray), both expression-based approaches. To ensure comparability, all models were trained, validated, and tested using identical patient cohorts for each cancer–drug pair. The ten pairs include solid tumor types (BLCA, CESC, COAD, LGG, LUAD, and STAD) treated with corresponding anti-cancer agents. (**C**) Distribution of AUROC values across the ten cancer–drug pairs. Each point represents one pair. Box plots display the interquartile range (IQR), with the median indicated by the center line and whiskers extending to 1.5×IQR. Statistical significance between SubNetDL and each model was assessed using paired two-sided Wilcoxon signed-rank tests, with *P*-values shown above the brackets.

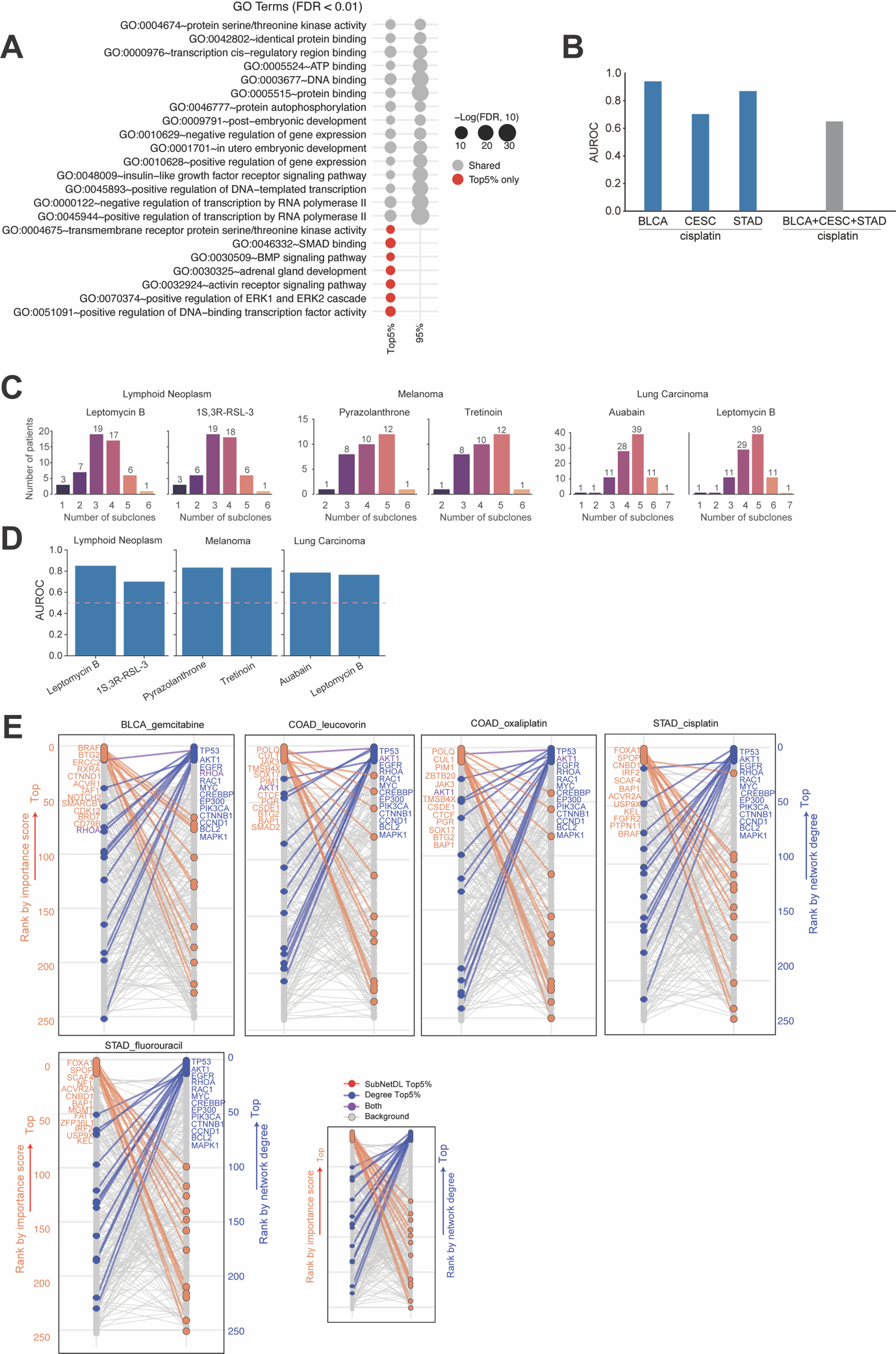

**Supplementary Figure 4.** Biological characterization and independent validation of SubNetDL predictions**, related to Figure 2 and 5**. (**A**) Gene ontology (GO) enrichment analysis for biological process (BP) and molecular function (MF) was performed on two groups: (1) genes that appeared in the top 5% of SubNetDL importance scores in at least one of the ten cancer type–drug combinations, and (2) genes that were never ranked in the top 5% across any of the combinations. All human protein-coding genes were used as the background. (**B**) Comparison of cancer-type–specific and pooled pan-cancer models for cisplatin response prediction. AUROC values of SubNetDL trained on cisplatin-treated patients are shown for three cancer-type–specific models—bladder urothelial carcinoma (BLCA), cervical squamous cell carcinoma (CESC), and stomach adenocarcinoma (STAD) (blue bars)—and for a pooled pan-cancer model trained on the combined cohort (BLCA+CESC+STAD; gray bar). While cancer-type–specific models achieved high predictive performance (median AUROC = 0.87), the pooled pan-cancer model showed substantially reduced performance (AUROC = 0.65), indicating that cancer-type–specific patterns contribute significantly to prediction accuracy. (**C**) Inferred subclonal architecture in CCLE cell lines. Histograms show the number of mutation-derived subclones per cell line across selected cancer types (Lymphoid Neoplasm, Melanoma, and Lung Carcinoma) and drug treatments (e.g., Leptomycin B, Tretinoin, Ouabain). Subclonal structures were inferred using the same clustering framework applied to TCGA samples. A median of two subclones per cell line was detected. (**D**) Predictive performance in independent preclinical models. AUROC values are shown for SubNetDL predictions across CCLE cell line–drug pairs. Drug response was defined using IC50 measurements from the Genomics of Drug Sensitivity in Cancer (GDSC) database, with cell lines stratified into responders (lower 50% IC50) and non-responders (upper 50% IC50). SubNetDL achieved a median AUROC of 0.81 (range: 0.70–0.85). (**E**) Genes ranked in the top 5% by SubNetDL are labeled in red, those ranked in the top 5% by network degree (based on the STRING PPI network) are labeled in blue, and genes ranked in the top 5% by both are shown in purple. All other genes are shown in grey.

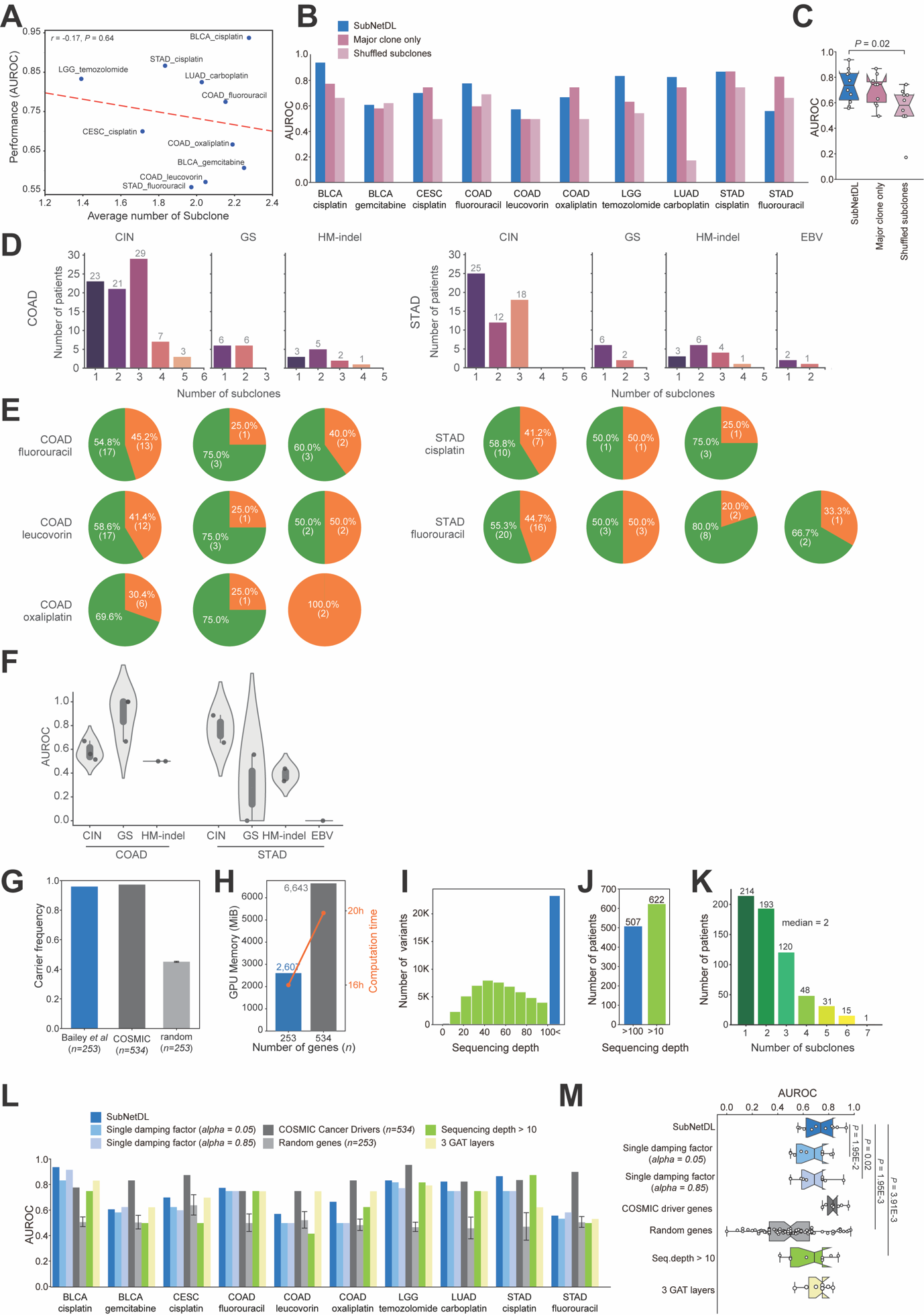

**Supplementary Figure 5.** Subclonal heterogeneity–associated predictive signals and robustness analyses of the SubNetDL framework**, related to Figure 2**. (**A**) Association between subclonal heterogeneity and predictive performance across cancer types. The average number of subclones detected per cancer type was compared with the corresponding prediction performance (AUROC) across ten cancer type–drug combinations. Each point represents a single cancer type–treatment pair. Pearson’s correlation was calculated between the average subclone count and AUROC values across cancer types. (**B**) Assessment of model dependence on subclonal organization. AUROC values of SubNetDL (blue) are compared across ten cancer–drug cohorts against two perturbed versions: restricting input to the major clone only (dark pink) and randomly shuffling mutations across inferred subclones (light pink). (**C**) Distribution of predictive performance under perturbations. Box plots summarize the AUROC distributions across the evaluated cohorts for each configuration. Randomly shuffling mutations resulted in a statistically significant decrease in performance (median AUROC 0.73 vs. 0.58; *P* = 0.02), while the major-clone-only model maintained comparable performance to the full subclonal model (median AUROC 0.74 vs. 0.73; *P* = 0.43). Center lines indicate medians, boxes represent interquartile ranges (IQR), and whiskers extend to 1.5×IQR. Statistical differences were assessed using a two-sided Wilcoxon signed-rank test with *P*-values shown above the brackets. Only statistically significant differences (*P* < 0.05) are indicated in the figure. (**D**) Distribution of subclone counts by molecular subtype. Bar plots show the number of patients stratified by the detected number of subclones in COAD (left) and STAD (right) across molecular subtypes: CIN (chromosomal instability), GS (genomically stable), HM-indel (hypermutated), and EBV (Epstein–Barr virus). Numbers above bars indicate the absolute number of patients in each subclone category. (**E**) Treatment response rates by molecular subtype. Pie charts depict the proportion of responders (green) and non-responders (orange) to common therapeutic agents (fluorouracil, leucovorin, oxaliplatin, and cisplatin). Percentages and absolute patient numbers are shown within each segment. (**F**) Predictive performance of the model across molecular subtypes. Box plots show the interquartile range (IQR), with center lines indicating the median and whiskers extending to 1.5×IQR. (**G**) Proportion of patients harboring at least one mutation within different driver gene sets. The curated Bailey *et al*. set (*n* = 253 genes) captures > 95.9% of mutation carriers across analyzed cohorts. (**H**) Computational cost at different input gene scales. The bars represent the measured peak GPU memory consumption (MiB), and the orange line indicates total training time for models trained with *n* = 253 and 534 genes. (**I**) Distribution of sequencing depth across somatic variants included in subclonal inference. Variants meeting the original depth criterion (>100, blue) are shown alongside additional variants included under the relaxed criterion (10–100, green). (**J**) Comparison of evaluable patient counts under stringent (>100) versus relaxed (>10) sequencing depth thresholds, showing an increase from 507 to 622 patients with relaxed filtering. (**K**) Distribution of inferred subclone counts per patient (sequencing depth >10). (**L**) AUROC comparison across ten cancer–drug pairs for the baseline SubNetDL (dark blue) and alternative configurations: single damping factor models (α = 0.05, steel blue; α = 0.85, gray-blue), expanded driver set using COSMIC genes (dark gray), random gene backbones (light gray), models trained with relaxed variant filtering (light green) and models in which the number of GAT layers was increased from the baseline of two to three (yellow). (**M**) Summary box plots of AUROC distributions for each configuration. Statistical comparisons were performed against the baseline SubNetDL using paired two-sided Wilcoxon signed-rank tests. Only statistically significant differences (*P* < 0.05) are indicated in the figure.
